# Functional lability of RNA-dependent RNA polymerases in animals

**DOI:** 10.1101/339820

**Authors:** Natalia Pinzón, Stéphanie Bertrand, Lucie Subirana, Isabelle Busseau, Hector Escrivá, Hervé Seitz

## Abstract

RNA interference (RNAi) requires RNA-dependent RNA polymerases (RdRPs) in many eukaryotes, and RNAi amplification constitutes the only known function for eukaryotic RdRPs. Yet in animals, classical model organisms can elicit RNAi without possessing RdRPs, and only nematode RNAi was shown to require RdRPs. Here we show that RdRP genes are much more common in animals than previously thought, even in insects, where they had been assumed not to exist. RdRP genes were present in the ancestors of numerous clades, and they were subsequently lost at a high frequency. In order to probe the function of RdRPs in a deuterostome (the cephalochordate *Branchiostoma lanceolatum*), we performed high-throughput analyses of small RNAs from various *Branchiostoma* developmental stages. Our results show that *Branchiostoma* RdRPs do not appear to participate in RNAi: we did not detect any candidate small RNA population exhibiting classical siRNA length or sequence features. Our results show that RdRPs have been independently lost in dozens of animal clades, and even in a clade where they have been conserved (cephalochordates) their function in RNAi amplification is not preserved. Such a dramatic functional variability reveals an unexpected plasticity in RNA silencing pathways.

**Author summary:** RNA interference (RNAi) is a conserved gene regulation system in eukaryotes. In non-animal eukaryotes, it necessitates RNA-dependent RNA polymerases (”RdRPs”). Among animals, only nematodes appear to require RdRPs for RNAi. Yet additional animal clades have RdRPs and it is assumed that they participate in RNAi. Here, we find that RdRPs are much more common in animals than previously thought, but their genes were independently lost in many lineages. Focusing on a species with RdRP genes (a cephalochordate), we found that it does not use them for RNAi. While RNAi is the only known function for eukaryotic RdRPs, our results suggest additional roles. Eukaryotic RdRPs thus have a complex evolutionary history in animals, with frequent independent losses and apparent functional diversification.

## Introduction

Small interfering RNAs (siRNAs) play a central role in the RNA interference (RNAi) response. Usually loaded on a protein of the AGO subfamily of the Argonaute family, they recognize specific target RNAs by sequence complementarity and typically trigger their degradation by the AGO protein [1]. In many eukaryotic species, normal siRNA accumulation requires an RNA-dependent RNA polymerase (RdRP). For example in plants, RdRPs are recruited to specific template RNAs and they generate long complementary RNAs [2–4]. The template RNA and the RdRP product are believed to hybridize, forming a long double-stranded RNA which is subsequently cleaved by Dicer nucleases into double-stranded siRNAs (reviewed in [5]). In fungi, RdRPs have also been implicated in RNAi and in RNA-directed heterochromatinization [6–9], but the exact nature of their products remains elusive: fungal RdRPs are frequently proposed to polymerize long RNAs which can form Dicer substrates after annealing to the RdRP template [10–12]. But the purified *Neurospora crassa, Thielavia terrestris* and *Myceliophthora thermophila* QDE-1 RdRPs tend to polymerize essentially short (9–21 nt) RNAs *in vitro*, suggesting that they may generate Dicer-independent small RNAs [13, 14]. In various unicellular eukaryotes, RdRPs have also been implicated in RNAi and related mechanisms (*e.g.*, see [15, 16]). It is usually believed that their products are long RNAs that anneal with the template to generate a Dicer substrate, and that model has gained experimental support in one organism, *Tetrahymena* [17].

Among eukaryotes, animals are thought to constitute an exception: most classical animal model organisms (*Drosophila* and mammals) can elicit RNAi without the involvement of an RdRP [1]. Only one animal model organism was shown to require RdRPs for RNAi: the nematode *Cænorhabditis elegans* [18, 19]. In nematodes, siRNAs made by Dicer only constitute a minor fraction of the total siRNA pool: such “primary” siRNAs recruit an RdRP on target RNAs, triggering the production of short antisense RNAs named “secondary siRNAs” [20–22]. Secondary siRNAs outnumber primary *≈*siRNAs by 100-fold [20] and the major class of secondary siRNAs (the so-called “22G RNAs”) is loaded on proteins of the WAGO subfamily of the Argonaute family [22, 23]. WAGO proteins appear to be unable to cleave RNA targets [23]. Yet WAGO/secondary siRNA/cofactor complexes appear to be much more efficient at repressing mRNA targets than AGO/primary siRNA/cofactor complexes [24], possibly by recruiting another, unknown, nuclease. In contrast to Dicer products (which bear a 5’ monophosphate), direct RdRP products bear a 5’ triphosphate. 22G RNAs are thus triphosphorylated on their 5’ ends [20]. Another class of nematode RdRP products, the “26G RNAs”, appears to bear a 5’ monophosphate, and it is not clear whether they are matured from triphosphorylated precursors, or whether they are directly produced as monophosphorylated RNAs [25–27].

The enzymatic activity of RNA-dependent RNA polymerization can be mediated by several unrelated protein families [28]. Most of these families are specific to viruses (*e.g.*, PFAM ID #PF00680, PF04196 and PF00978). Viral RdRPs are involved in genome replication and transcription in RNA viruses, and they share common structural motifs [29]. On the other hand, RdRPs involved in RNAi in plants, fungi and nematodes belong to a family named “eukaryotic RdRPs” (PFAM ID #PF05183). While viral RdRPs are conceivably frequently acquired by virus-mediated horizontal transfer, members of the eukaryotic RdRP family are thought to be inherited vertically only [30]. The eukaryotic RdRP family can be further divided into three subfamilies, named *α, β* and *γ* based on sequence similarity. Phylogenetic analyses suggest these three subfamilies derive from three ancestral RdRPs that could have coexisted in the most recent common ancestor of animals, fungi and plants [31].

Besides eukaryotic RdRPs, other types of RdRP enzymes have been proposed to exist in various animals. It has been suggested that human cells express an atypical RdRP, composed of the catalytic subunit of telomerase and a non-coding RNA [32]. While that complex exhibits RdRP activity *in vitro*, functional relevance of that activity is unclear, and other mammalian cells were shown to perform RNAi without RdRP activity [33]. More recently, bat species of the *Eptesicus* clade were shown to possess an RdRP of viral origin, probably acquired upon endogenization of a viral gene at least 11.8 million years ago [34].

Here we took advantage of the availability of hundreds of metazoan genomes to draw a detailed map of predicted RdRP genes in animals. We found RdRP genes in a large diversity of animal clades, even in insects, where they had escaped detection so far. Even though RdRP genes are found in diverse animal clades, they are lacking in many species, indicating that they were frequently and independently lost in many lineages. Furthermore, the presence of RdRP genes in non-nematode genomes raises the possibility that additional metazoan lineages possess an RdRP-based siRNA amplification mechanism. We sequenced small RNAs from various developmental stages in one such species with 6 candidate RdRP genes, the cephalochordate *Branchiostoma lanceolatum*, using experimental procedures that were designed to detect both 5’ mono- and tri-phosphorylated RNAs. Our analyses did not reveal any evidence of the existence of secondary siRNAs in that organism. While RNAi is the only known function for eukaryotic RdRPs, we thus propose that *Branchiostoma* RdRPs do not participate in RNAi.

## Materials and methods

### Bioinformatic analyses of protein sequences

Predicted animal proteome sequences were downloaded from the following databases:

NCBI (ftp://ftp.ncbi.nlm.nih.gov/genomes/), VectorBase

(https://www.vectorbase.org/download/), FlyBase

(ftp://ftp.flybase.net/releases/FB2015_03/), JGI

(ftp://ftp.jgi-psf.org/pub/JGI_data/), Ensembl

(ftp://ftp.ensembl.org/pub/release-81/fasta/), WormBase

(ftp://ftp.wormbase.org/pub/wormbase/species/) and Uniprot

(http://www.uniprot.org/). The predicted *Branchiostoma lanceolatum* proteome was obtained from the *B. lanceolatum* genome consortium. RdRP HMMer profiles were downloaded from PFAM v. 31.0 (http://pfam.xfam.org/): 19 viral RdRP family profiles (PF00602, PF00603, PF00604, PF00680, PF00946, PF00972, PF00978, PF00998, PF02123, PF03035, PF03431, PF04196, PF04197, PF05788, PF05919, PF07925, PF08467, PF12426, PF17501) and 1 eukaryotic RdRP family profile (PF05183). Candidate RdRPs were selected by **hmmsearch** with an E-value cutoff of 10*-*^2^. Only those candidates with a complete RdRP domain according to NCBI’s Conserved domain search tool (https://www.ncbi.nlm.nih.gov/Structure/bwrpsb/bwrpsb.cgi) were considered (tolerating up to 20% truncation on either end of the domain). One identified candidate, in the bat *Rhinolophus sinicus*, appears to be a plant contaminant (it is most similar to plant RdRPs, and its genomic scaffold [ACC# LVEH01002863.1] only contains that gene): it was not included in Figure 1 and in Supplementary Fig. S1.

**Fig 1.**
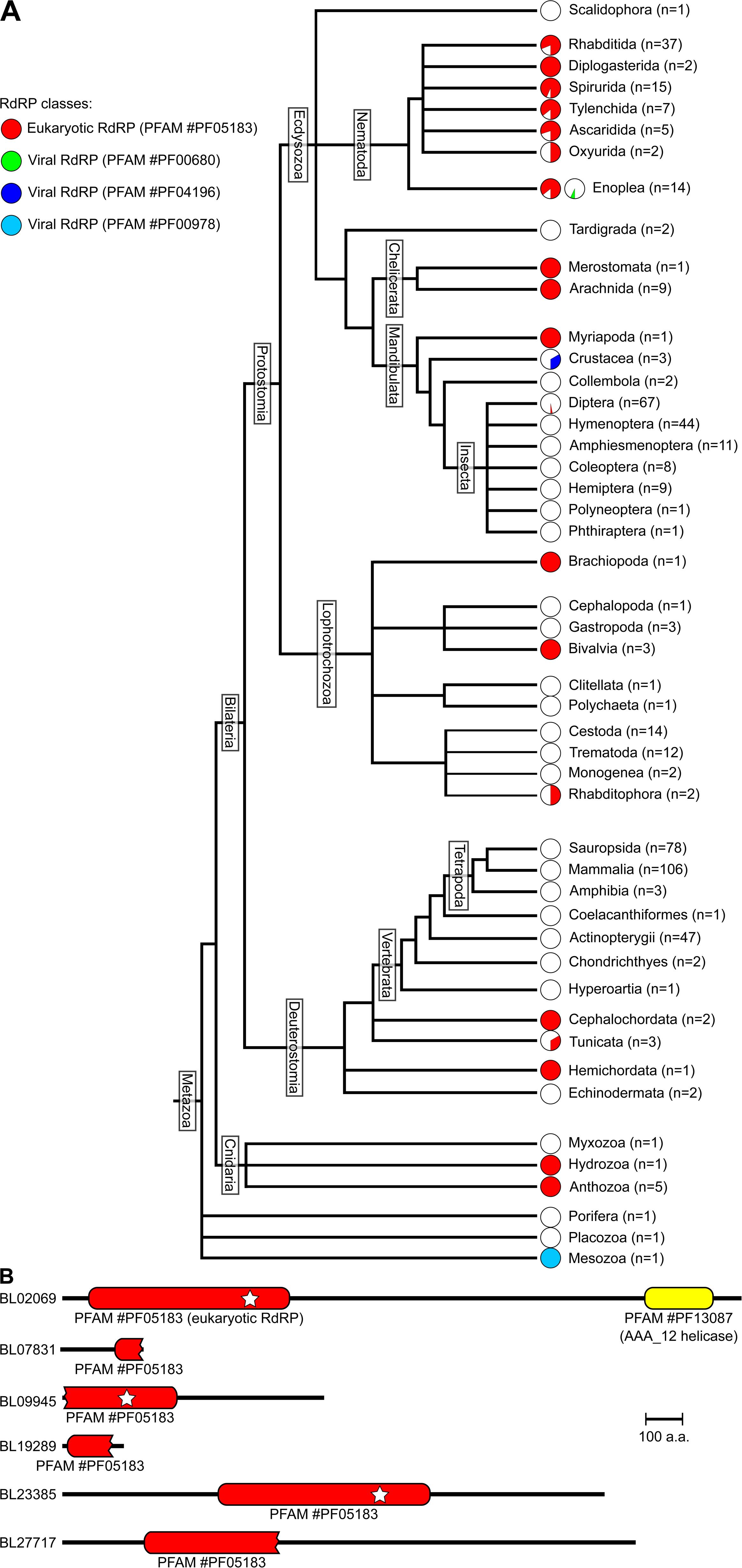
Phylogenetic distribution of RdRP genes in metazoans. A. Proteome sequences from 538 metazoans were screened for potential RdRPs. For each clade indicated on the right edge, *n* is the number of species analyzed in the clade, and piecharts indicate the proportion of species possessing RdRP genes (with each RdRP family represented by one piechart, according to the color code given at the top left). B. An HMMer search identifies 6 candidate RdRPs in the predicted *Branchiostoma lanceolatum* proteome. Only 2 candidates have a complete RdRP domain (represented by a red bar with round ends; note that apparent domain truncations may be due to defective proteome prediction). A white star indicates that every catalytic amino acid is present. Candidate BL02069 also possesses an additional known domain, AAA 12 (in yellow).

The *Branchiostoma* Hen1 candidate was identified using **HMMer** on the predicted *lanceolatum* proteome, with an HMMer profile built on an alignment of *Drosophila melanogaster, Mus musculus, Danio rerio, Nematostella vectensis* and *Arabidopsis thaliana* Hen1 sequences.

### Phylogenetic tree reconstruction

Amino acid sequences of the eukaryotic RdRP domain (Pfam #PF05183) were retrieved from PFAM [35], and supplemented with the RdRP domains of the proteins identified in the 538 animal proteomes (*cf* above). Sequences were aligned using **hmmalign** [36] using the HMM profile of the PF05183 RdRP domain. Sequences for which the domain was incomplete were deteled from the alignment. Sites used to reconstruct the phylogenetic tree were selected using **trimAl** [37] on the Phylemon 2.0 webserver [38]. Bayesian inference (BI) tree was inferred using **MrBayes 3.2.6** [39], with the model recommended by **ProtTest 1.4** [40] under the Akaike information criterion (LG+Γ), at the CIPRES Science Gateway portal [41]. Two independent runs were performed, each with 4 chains and one million generations. A burn-in of 25% was used and a fifty majority-rule consensus tree was calculated for the remaining trees. The obtained tree was customized using **FigTree v.1.4.0**.

### Sample collection

Mediterranean amphioxus (*Branchiostoma lanceolatum*) males and females were collected at le Racou (Argelès-sur-mer, France) and were induced to spawn as previously described [42]. Embryos were obtained after fertilization in Petri dishes filled with filtered sea water and cultivated at 19*°*C. Total RNA was extracted from 8, 15, 36 and 60 hours post fertilization (hpf) embryos (three independent batches for each stage, pooled before small RNA gel purification) as well as from males (6 pooled individuals) and females (4 pooled individuals) using the RNeasy mini kit (for embryonic samples) and the RNeasy midi kit (for adult samples) (Qiagen).

### Sequencing analyses

The BL09945 locus was PCR-amplified from adult female DNA, cloned in the pGEM-T easy vector (cat. #A1360; Promega, Madison, WI, USA) and sequenced by MWG Eurofins Genomics (Ebersberg, Germany).

For Small RNA-Seq, 18–30 nt RNAs were gel-purified from total RNA (using between 92 and 228 *µ*g total RNA per sample). One quarter of the small RNA preparation was kept untreated before library preparation (for “Libraries #1”). One quarter was incubated for 10 min at room temperature in 100 *µ*L of freshly-prepared 60 mM sodium borate (pH=8.6), 25 mM sodium periodate, then the reaction was quenched with 10 *µ*L glycerol (for “Libraries #2”). One quarter was treated with 1.25 U Terminator exonuclease (Epicentre, Madison, WI, USA) in 25 *µ*L 1X Terminator reaction buffer A for 1h at 30*°*C, then the reaction was quenched with 1.25 *µ*L 500 mM EDTA (pH=8.0) and ethanol-precipitated. RNA was then treated with 5 U Antarctic phosphatase (New England Biolabs, Ipswich, MA, USA) in 20 *µ*L 1X Antarctic phosphatase buffer for 30 min at 37*°*C, the enzyme was heat-inactivated, then RNA was precipitated, then phosphorylated by 15 U T4 PNK (New England Biolabs) with 50 nmol ATP in 50 *µ*L 1X T4 PNK buffer for 30 min at 37*°*C, then the enzyme was heat-inactivated (for “Libraries #3”). One quarter was treated successively with Terminator exonuclease, Antarctic phosphatase, T4 PNK then boric acid and sodium periodate, with the same protocols (for “Libraries #4”). Small RNA-Seq libraries were then generated using the TruSeq Small RNA library preparation kit (Illumina, San Diego, CA, USA), following the manufacturer’s instructions.

Libraries were sequenced by the MGX sequencing facility (CNRS, Montpellier, France). Read sequences were aligned on the *B. lanceolatum* genome assembly [43] using **bowtie2**. A database of abundant non-coding RNAs was assembled by a search for orthologs for human and murine rRNAs, tRNAs, snRNAs, snoRNAs and scaRNAs; deep-sequencing libraries were also mapped on that database using **bowtie2**, and matching reads were flagged as “abundant ncRNA fragments”. For pre-miRNA annotation, every *B. lanceolatum* locus with a Blast E-value ≤ 10*-*^6^ to any of the annotated *B. floridae* or *B. belcheri* pre-miRNA hairpins in miRBase v.22 was selected. Reads matching these loci were identified using **bowtie2**. For the measurement of miRNA abundance during development, hairpins were further screened for their **RNAfold**-predicted secondary structure and their read coverage: Supplementary Table S1 only lists unbranched hairpins with at least 25 bp in their stem, with a predicted Δ*G*_*folding*_ *≤ -* 15 *kcal.mol*^*-*1^, generating mostly 21- to 23-mer RNAs, and with at least 20 ppm read coverage on any nucleotide of the hairpin.

RNA-Seq data was taken in [43] for embryonic and juvenile samples. Adult sample libraries were prepared and sequenced by “Grand plateau technique r’egional de g’enotypage” (SupAgro-INRA, Montpellier). mRNA abundance data was extracted using **vast-tools** [44].

### Extragenomic contig assembly and annotation

Small RNA reads that fail to map on the *B. lanceolatum* genome or transcriptome according to **bowtie2** were collected and assembled using **velvet** [45], with *k* values ranging from 9 to 19 for better sensitivity [46].

Contigs at least 50 bp in length were then compared to the NCBI non-redundant nucleotide collection (as of October 31, 2018) by **megablast** on the NCBI server with default parameters. Contigs with a detected similarity to known sequences in the collection were annotated with phylogenetic information (see Table 2) using the NCBI “Taxonomy” database.

### Code availability

Source code, detailed instructions, and intermediary data files are accessible on GitHub (https://github.com/HKeyHKey/Pinzon_et_al_2018) as well as on https://www.igh.cnrs.fr/en/research/departments/genetics-development/systemic-impact-of-small-regulatory-rnas/165-computer-programs.

### Data availability

Deep-sequencing data has been deposited at NCBI’s Short Read Archive under BioProject accession #PRJNA419760 (for Small RNA-Seq) and BioSample accession #SAMN09381006 and SAMN09381007 (for adult RNA-Seq). Sequences of the re-sequenced *B. lanceolatum* BL09945 locus have been deposited at GenBank under accession #MH261373 and #MH261374.

## Results

### A sporadic phylogenetic distribution of RdRP genes

Previous analyses showed that a few animal genomes contain candidate RdRP genes [28, 31, 34, 47]. Rapid development of sequencing methods recently made many animal genomes available, allowing a more complete coverage of the phylogenetic tree. A systematic search for RdRP candidates (including every known viral or eukaryotic RdRP family) in 538 predicted metazoan proteomes confirms that animal species possessing RdRPs are unevenly scattered in the phylogenetic tree, but they are much more abundant than previously thought: we identified 98 metazoan species with convincing eukaryotic RdRP genes (see Figure 1A). Most RdRPs identified in animal predicted proteomes belong to the eukaryotic RdRP family, but 3 species (the Enoplea *Trichinella murrelli*, the Crustacea *Daphnia magna* and the Mesozoa *Intoshia linei)* possess RdRP genes belonging to various viral RdRP families (in green, dark blue and light blue on Figure 1A), which were probably acquired by horizontal transfer from viruses. Most sequenced nematode species appear to possess RdRP genes. But in addition, many other animal species are equipped with eukaryotic RdRP genes, even among insects (the Diptera *Clunio marinus* and *Rhagoletis zephyria*), where RdRPs were believed to be absent [47, 48].

Our observation of eukaryotic family RdRPs in numerous animal clades therefore prompted us to revisit the evolutionary history of animal RdRPs: eukaryotic RdRPs were probably present in the last ancestors for many animal clades (including insects, mollusks, deuterostomes) and they were subsequently lost independently in most insects, mollusks and deuterostomes. It has been recently shown that the last ancestor of arthropods possessed an RdRP, which was subsequently lost in some lineages [47]: that result appears to be generalizable to a large diversity of animal clades. The apparent absence of RdRPs in some species may be due to genome incompleteness, or to defective proteome prediction. Excluding species with low numbers of long predicted proteins (⩾500 or 1,000 amino acids) indeed eliminates a few dubious proteomes, but the resulting distribution of RdRPs in the phylogenetic tree is only marginally affected, and still suggests multiple recent RdRP losses in diverse lineages (see Supplementary Fig. S1).

Alternatively to multiple gene losses, such a sporadic phylogenetic distribution could be due to frequent horizontal transfer of RdRP genes in animals. In order to assess these two possibilities, it is important to better understand the evolution of metazoan RdRPs in the context of the whole eukaryotic RdRP family. We therefore used sequences found in all eukaryotic groups for phylogenetic tree reconstruction. The supports for deep branching are low and do not allow us to propose a complete evolutionary history scenario of the whole eukaryotic RdRP family (see Figure 2A). However, metazoan sequences are forming three different groups, which were named RdRP *α, β* and *γ* according to the pre-existing nomenclature [31], and their position in relation to non-metazoan eukaryotic sequences does not support an origin through horizontal gene transfer. The only data that would support horizontal gene transfer pertains to the metazoan sequences of the RdRP *β* group (see Figure 2C). Indeed, sequences of stramenopiles and a fungus belonging to parasitic species are embedded in this clade. For the RdRP *α* and *γ* groups, the phylogeny strongly suggests that they derive from at least two genes already present in the common ancestor of cnidarians and bilaterians and that the scarcity of RdRP presence in metazoans would be the result of many secondary gene losses. Even the *Strigamia maritima* RdRP was probably not acquired by a recent horizontal transfer from a fungus, as has been proposed [47]: when assessed against a large number of eukaryotic RdRPs, the *S. maritima* sequence clearly clusters within metazoan *γ* RdRP sequences. In summary, we conclude that RdRPs were present in the last ancestors of many animal clades, and they were recently lost independently in diverse lineages.

**Fig 2.**
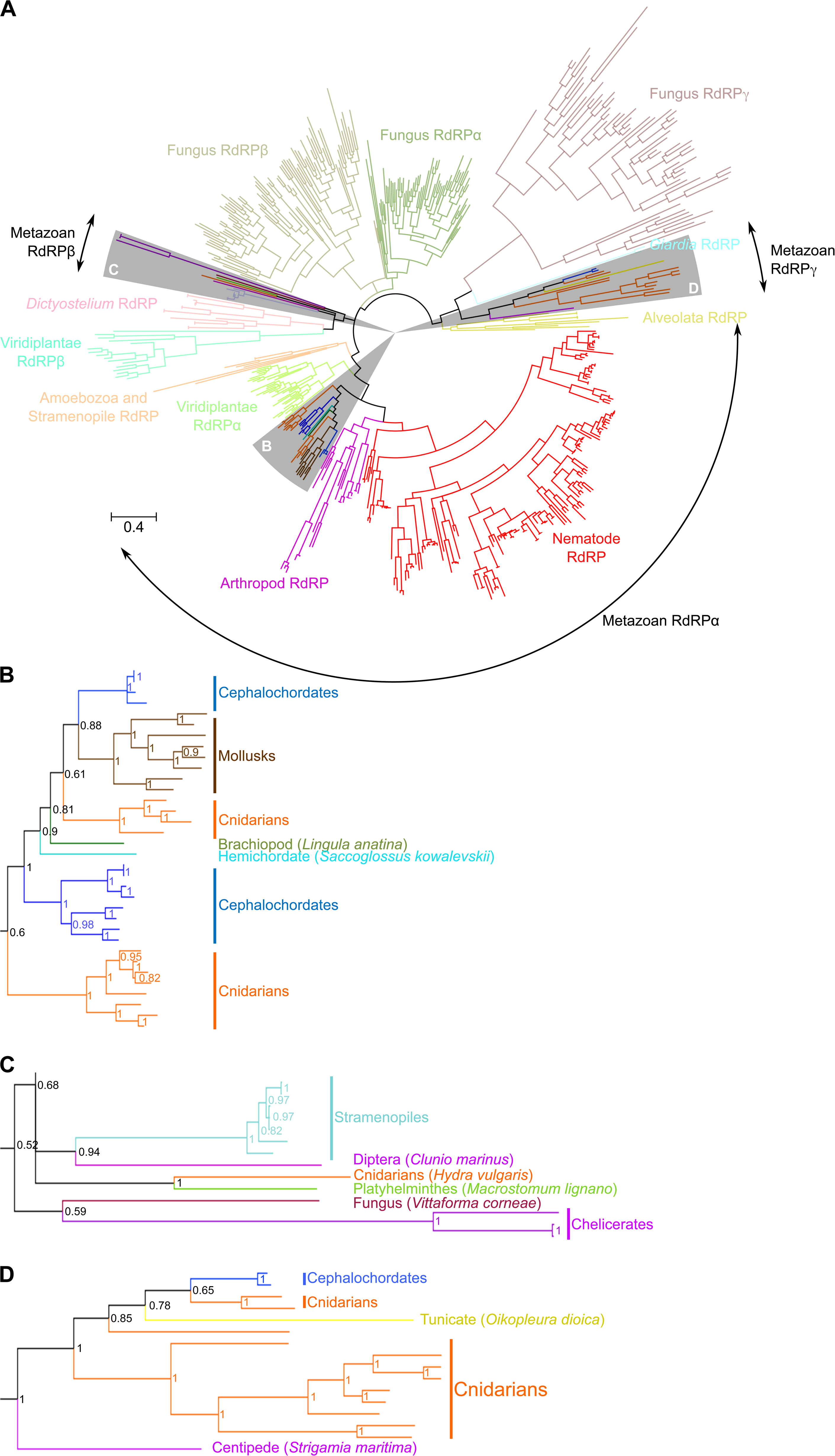
Eukaryotic RdRP phylogeny supports the vertical transfer scenario. Bayesian phylogenetic tree of the eukaryotic RdRP family. *α, β* and *γ* clades of eukaryotic RdRPs have been defined by [31]. Sectors highlighted in grey are detailed in panels B, C and D for clarity. Scale bar: 0.4 amino acid substitution per position. Posterior probability values are indicated for each node in panels B–D.

### Experimental search for RdRP products in *Branchiostoma*

In an attempt to probe the functional conservation of RdRP-mediated RNAi amplification among metazoans, we decided to search for secondary siRNAs in an organism where RdRP candidates could be found, while being distantly related to *C. elegans*. We reasoned that endogenous RNAi may act as a gene regulator during development or as an anti-pathogen response. Thus siRNAs are more likely to be detected if several developmental stages are probed, and if the analyzed specimens are gathered in a natural ecosystem, where they are naturally challenged by pathogens. From these considerations it appears that the most appropriate organism is a cephalochordate species, *Branchiostoma lanceolatum* [49]. In good agreement with the known scarcity of gene loss in that lineage [50], cephalochordates also constitute the only bilaterian clade for which both RdRP *α* and *γ* sequences can be found, thus increasing the chances of observing RNAi amplification despite the diversification of eukaryotic RdRPs into three groups. According to our HMMer-based search, the *B. lanceolatum* genome encodes 6 candidate RdRPs, three of which containing an intact active site DbDGD (with b representing a bulky amino acid; [51]) (see Figure 1B). The current *B. lanceolatum* genome assembly contains a direct 1,657 bp repeat in one of the 6 RdRP genes, named BL09945. This long duplication appears to be an assembly artifact: we cloned and re-sequenced that locus and identified two alleles (with a synonymous mutation on the 505th codon; deposited at GenBank under accession numbers MH261373 and MH261374), and none of them contained the repeat. In subsequent analyses, we thus used a corrected version of that locus, where the 1,657 bp duplication is removed.

In most metazoan species, siRNAs (as well as miRNAs) bear a 5’ monophosphate and a 3’ hydroxyl [52, 53]. The only known exceptions are “22G” secondary siRNAs in nematodes (they bear a 5’ triphosphate; [20]), which may be primary polymerization products by an RdRP; Ago2-loaded siRNAs and miRNA in *Drosophila*, which are 3’-methylated on their 2’ oxygen after loading on Ago2 and unwinding [54, 55]; and a subset of “26G” secondary siRNAs in nematodes (those which are loaded on the ERGO-1 Argonaute protein), which also bear a 2’-*O* -methyl on their 3’ end [56–58].

In order to detect small RNAs with any number of 5’ phosphates, bearing either an unmodified or a methylated 3’ end, we prepared multiple Small RNA-Seq libraries (see Figure 3A). Total RNA was extracted from various embryonic stages: gastrula (8 hours post-fertilization, hpf), early neurula (15 hpf), premouth neurula (36 hpf) and larvae (60 hpf), as well as from adult male and female specimens collected from their natural ecosystem. Small (18 to 30 nt long) RNAs were gel-purified, then Small RNA-Seq libraries were prepared using either the standard Small RNA-Seq protocol (which detects 5’ monophosphorylated small RNAs, whether they bear a 3’ methylation or not; “Library #1”); or by oxidizing small RNAs with NaIO4 in the presence of H3BO3 prior to library preparation (such treatment renders unmodified 3’ RNAs non-ligatable, hence undetectable by deep-sequencing; [59]; “Library #2”); or by treating small RNAs with the Terminator exonuclease (which degrades 5’ monophosphorylated RNAs) then with phosphatase then T4 PNK (to convert 5’ polyphosphorylated RNAs and 5’ hydroxyl RNAs into monophosphorylated RNAs, suitable for Small RNA-Seq library preparation; “Library #3”); or by a combination of both treatments (to detect only small RNAs bearing a 5’ polyphosphate or a 5’ hydroxyl, and a 3’ modification; “Library #4”). If the same experiments were performed in classical animal model organisms, such as *Drosophila*, nematodes and vertebrates (where miRNAs are essentially 5’ monophosphorylated and 3’-unmodified, and piRNAs are 5’ monophosphorylated and 3’-methylated), miRNAs would be expected to be detected in Libraries #1 and piRNAs, in Libraries #1 and 2. Nematode “22G” siRNAs would be detected in Libraries #3.

**Fig 3.**
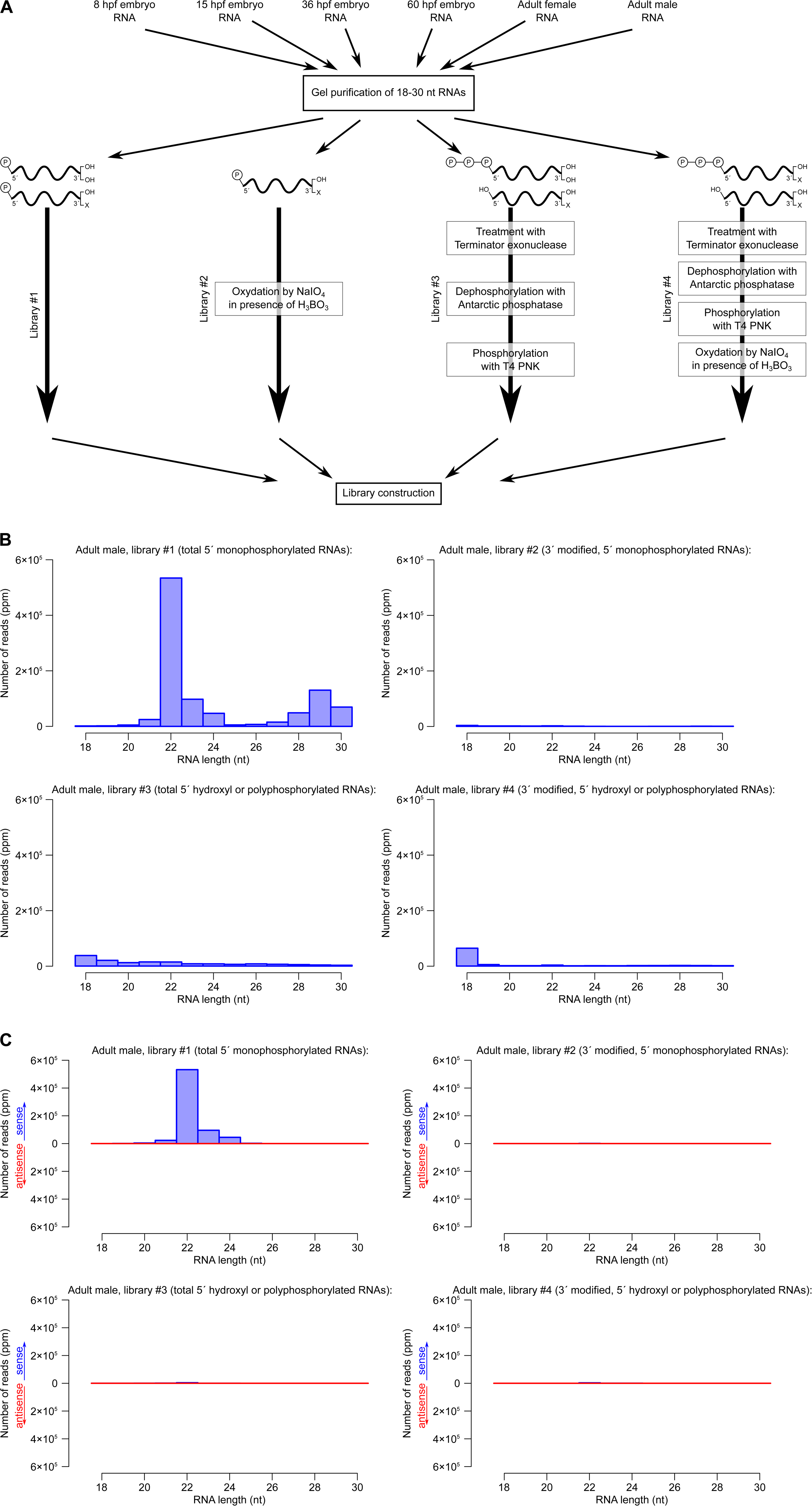
Detection of *B. lanceolatum* small RNAs. Four libraries were prepared for each biological sample, to detect small RNAs bearing either a single 5’ phosphate (Libraries #1 and 2) or any other number of phosphates (including zero; Libraries #3 and 4), and either a 2’-OH and 3’-OH 3’ end (Libraries #1 and 3), or a protected (*e.g.*, 2’-*O* -methylated) 3’ end (Libraries #2 and 4). hpf: hours post fertilization. B. Size distribution of genome-matching adult male small RNAs, excluding reads that match abundant non-coding RNAs (rRNAs, tRNAs, snRNAs, snoRNAs or scaRNAs). Read numbers are normalized by the total number of genome-matching reads (including *<*18 nt and *>*30 nt reads) that do not match abundant non-coding RNAs, and expressed as *parts per million* (ppm). C. Size distribution of adult male small RNAs matching pre-miRNA hairpins in the sense (blue) or antisense (red) orientation.

In the course of library preparation, it appeared that Libraries #4 contained very little ligated material, suggesting that small RNAs with a 3’ modification as well as *n* ⩾ 0 (with *n* ≠ 1) phosphates on their 5’ end, are very rare in *Branchiostoma* regardless of developmental stage. This observation was confirmed by the annotation of the sequenced reads: most reads in Libraries #4 did not map on the *B. lanceolatum* genome, probably resulting from contaminating nucleic acids (see Supplementary Fig. S2).

In Libraries #1 in each developmental stage, most *Branchiostoma* small RNA reads fall in the 18–30 nt range as expected. Other libraries tend to be heavily contaminated with shorter or longer reads, and 18–30 nt reads only constitute a small fraction of the sequenced RNAs (see Figure 3B for adult male libraries; see Supplementary File S1 section 1 for other developmental stages). miRNA loci have been annotated in two other cephalochordate species, *B. floridae* and *B. belcheri* (156 pre-miRNA hairpins for *B. floridae* and 118 for *B. belcheri* in miRBase v. 22). We identified the *B. lanceolatum* orthologous loci for annotated pre-miRNA hairpins from *B. floridae* or *B. belcheri*. Mapping our libraries on that database allowed us to identify candidate *B. lanceolatum* miRNAs. These RNAs are essentially detected in our Libraries #1, implying that, like in most other metazoans, *B. lanceolatum* miRNAs are mostly 22 nt long, they bear a 5’ monophosphate and no 3’ methylation (see Figure 3C for adult male libraries; see Supplementary File S1 section 2 for other developmental stages). Among the *B. lanceolatum* loci homologous to known *B. floridae* or *B. belcheri* pre-miRNA loci, 56 exhibit the classical secondary structure and small RNA coverage pattern of pre-miRNAs (*i.e.*, a stable unbranched hairpin generating mostly 21–23 nt long RNAs from its arms). These 56 loci, the sequences of the miRNAs they produce, and their expression profile during development, are shown in Supplementary Table S1.

### No evidence of RdRP-based siRNA amplification in *Branchiostoma*

In an attempt to detect siRNAs, we excluded every sense pre-miRNA-matching read and searched for distinctive siRNA features in the remaining small RNA populations. Whether RdRPs generate long antisense RNAs which anneal to sense RNAs to form a substrate for Dicer, or whether they polymerize directly short single-stranded RNAs which are loaded on an Argonaute protein, the involvement of RdRPs in RNAi should result in the accumulation of antisense small RNAs for specific target genes. These small RNAs should exhibit characteristic features:

- a narrow size distribution (imposed either by the geometry of the Dicer protein, or by the processivity of the RdRP [24, 60]; the length of Argonaute-loaded RNAs can also be further refined by exonucleolytic trimming of 3’ ends protruding from Argonaute [22, 61–65]);
- and possibly a sequence bias on their 5’ end; it is remarkable that the known classes of RdRP products in metazoans (nematode 22G and 26G RNAs) both display a strong bias for a guanidine at their 5’ end. RNA polymerases in general tend to initiate polymerization on a purine nucleotide [66–72] and it can be expected that primary RdRP products bear either a 5’A or a 5’G. Of note: loading on an Argonaute may also impose a constraint on the identity of the 5’ nucleotide, because of a sequence preference of either the Argonaute protein or its loading machinery [73–78].

The analysis of transcriptome-matching, non-pre-miRNA-matching small RNAs does not indicate that such small RNAs exist in *Branchiostoma* (see Figure 4 for adult males, and Supplementary File S1, section 3, for the complete data set). In early embryos, 5’ monophosphorylated small RNAs exhibit the typical size distribution and sequence biases of piRNA-rich samples: a heterogeneous class of 23 to 30 nt long RNAs. Most of them tend to bear a 5’ uridine, but 23 to 26 nt long RNAs in the sense orientation to annotated transcripts tend to have an adenosine at position 10 (especially when the matched transcript exhibits a long ORF; see Supplementary File S1, section 4). Vertebrate and *Drosophila* piRNAs display very similar size profiles and sequence biases [79–85]. These 23–30 nt long RNAs may thus constitute the *Branchiostoma* piRNAs, but surprisingly, they do not appear to bear a 2’-*O* -methylation on their 3’ end (see Discussion). Note that piRNAs appear to be mostly restricted to the germ line and gonadal somatic cells in other model organisms. But they are so abundant in piRNA-expressing cells, and so abundantly maternally deposited in fertilized eggs, that they can still be readily detected in embryonic or adult whole-body small RNA samples [25, 86–90]. It is thus not surprising to observe piRNA candidates in our *Branchiostoma* whole-body Small RNA-Seq libraries.

**Fig 4.**
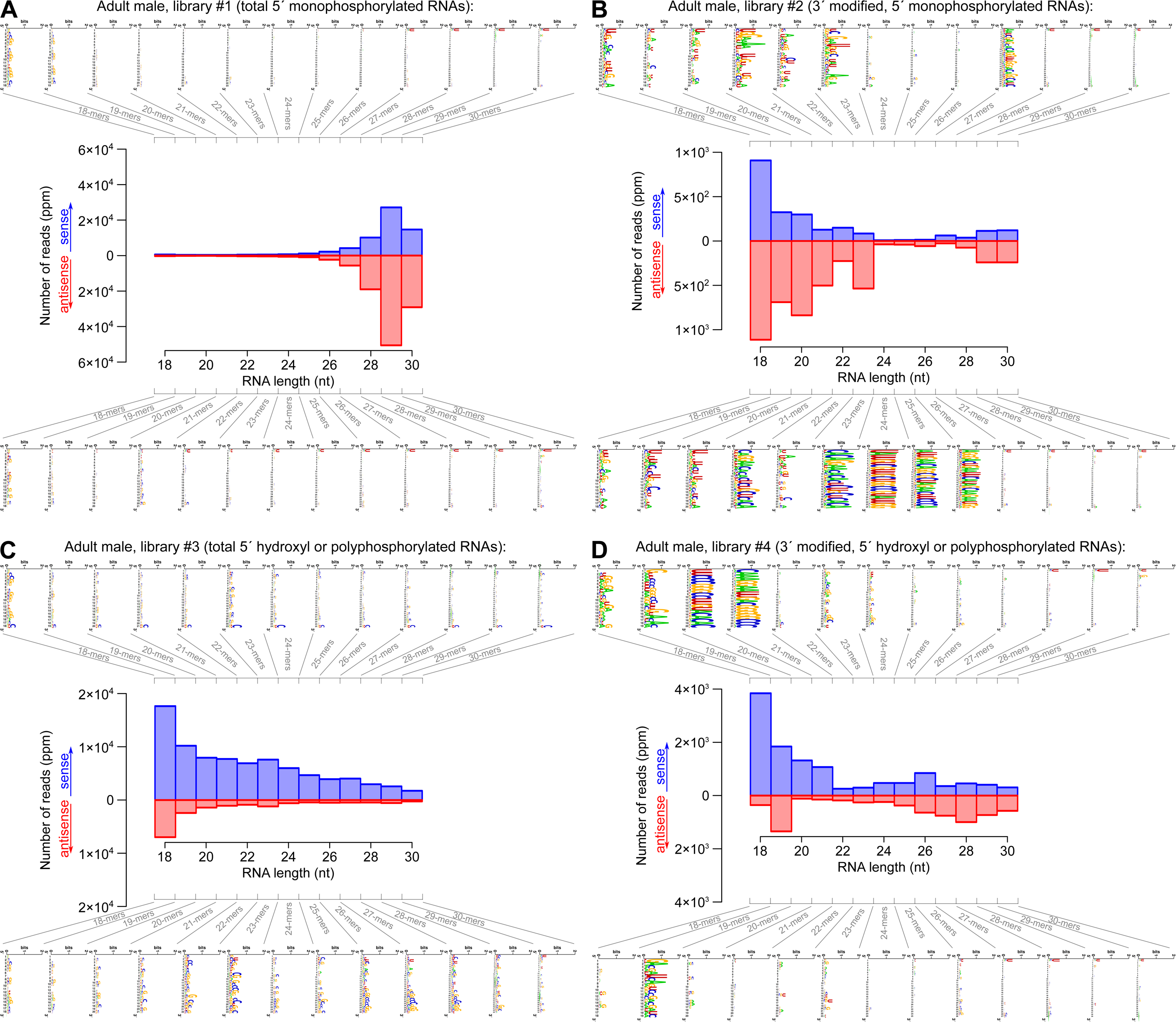
Size distribution and sequence logos for transcriptome-matching small RNAs in adult males. See Supplementary File S1, section 3, for the other developmental stages. A: Library #1, B: Library #2, C: Library #3, D: Library #4. Numbers of reads are expressed as *parts per million* (ppm) after normalization to the total number of genome-matching reads that do not match abundant non-coding RNAs. For each orientation (sense or antisense-transcriptome-matching reads), a logo analysis was performed on each size class (18 to 30 nt long RNAs).

In summary, transcriptome-matching small RNAs in our *Branchiostoma* libraries contain miRNA and piRNA candidates, but they do not contain any obvious class of presumptive secondary siRNAs that would exhibit a precise size distribution, and possibly a 5’ nucleotide bias. If *Branchiostoma* RdRPs generated secondary siRNAs by polymerizing mature short antisense RNAs (similarly to nematode 22G RNAs according to the prevalent model), then such hypothetical siRNAs should be detected in libraries #3. If *Branchiostoma* RdRPs generated long antisense RNAs, that would anneal to sense RNAs to produce a Dicer substrate (similarly to fungus and plant RdRP-derived siRNAs according to the prevalent model), then secondary siRNAs should be detected in libraries #1. As we did not observe candidate siRNA populations in either libraries #1 or 3, our data seem to rule out the existence of secondary siRNAs in *Branchiostoma*, regardless of the mechanistical involvement of RdRPs in their production.

One could imagine that transcriptome-matching siRNAs were missed in our analysis, because of issues with the *Branchiostoma* transcriptome assembly. It is also conceivable that siRNAs exist in *Branchiostoma*, but they do not match its genome or transcriptome (they could match pathogen genomes, for example if they contribute to an anti-viral immunity). We therefore analyzed other potential siRNA types: **(i)** genome-matching reads that do not match abundant non-coding RNAs (rRNAs, tRNAs, snRNAs, snoRNAs or scaRNAs); **(ii)** reads that match transcripts exhibiting long (100 codons, initiating on one of the three 5’-most AUG codons) open reading frames; **(iii)** reads that do not match the *Branchiostoma* genome, nor its transcriptome (potential siRNAs derived from pathogens). Once again, none of these analyses revealed any siRNA population in *Branchiostoma* (see detailed results in Supplementary File S1, sections 1, 4 and 5). This is in striking contrast to *Cænorhabditis elegans*, where antisense transcriptome-matching siRNAs (mostly 22 nt long, starting with a G) are easily detectable (see Supplementary File S1, section 6, for our analysis of publicly available *C. elegans* data; [22]).

### *Branchiostoma* RdRP activity is not clearly detected

Our failure to detect siRNA candidates may simply be due to the fact that they are poorly abundant in the analyzed developmental stages. In order to enrich for small RNA populations derived from RdRP activity, and exclude all the other types of small RNAs, we considered small RNAs mapping on exon-exon junctions in the antisense orientation. The antisense sequence of the splicing donor (GU) and acceptor (AG) sites does not constitute a donor/acceptor pair itself, implying that any RNA antisense to a spliced RNA must have originated from the action of an RdRP on the spliced RNA — it cannot derive from the splicing of an RNA transcribed in the antisense orientation.

We therefore selected all the 18–30 nt RNA reads that map on exon-exon junctions in the annotated transcriptome, and fail to map on the genome. Such reads map almost exclusively in the sense orientation (see Table 1). When focusing on the developmental stage where some transcripts exhibit the highest observed numbers of antisense exon-exon junction reads (15 hpf embryos, for the transcripts of genes BL05604 and BL00515), it appears that these antisense junction reads are highly homogeneous in sequence (sharing the same 5’ and 3’ ends), they do not map perfectly on the spliced transcript (with 1 mismatch in each), and their total abundance remains very small (less than 10 raw reads per transcript in a given developmental stage) (see Supplementary Fig. S3). RdRP genes themselves appear to be developmentally regulated, with candidate RdRPs harboring intact active sites showing expression peaks at 8 and 18 hpf (see Supplementary Fig. S4).

**Table 1.** Genes with highest read coverage on exon-exon junctions in the sense or antisense orientation.

| Whole transcriptome |  |  |  |  |  |
| --- | --- | --- | --- | --- | --- |
| Sense (ppm) |  |  | Antisense (ppm) |  |  |
| 94.2 |  |  | 1.33 |  |  |
| Top genes with sense-matching junction reads |  |  | Top genes with antisense-matching junction reads |  |  |
| Gene name | Sense reads (ppm) | Antisense reads (ppm) | Gene name | Sense reads (ppm) | Antisense reads (ppm) |
| BL01573 | 4.11 | 0 | BL05604 | 0 | 0.348 |
| BL21967 | 2.89 | 0 | BL00515 | 0.0332 | 0.149 |
| BL11086 | 2.29 | 0 | BL16381 | 0 | 0.0995 |
| BL01498 | 2.27 | 0 | BL06097 | 0.0663 | 0.0663 |
| BL23908 | 1.41 | 0 | BL13214 | 0 | 0.0497 |
| BL06958 | 1.38 | 0 | BL06086 | 0.0497 | 0.0497 |
| BL23502 | 1.18 | 0 | BL05692 | 0 | 0.0332 |
| BL20132 | 0.895 | 0 | BL11851 | 0.0332 | 0.0332 |
| BL10273 | 0.779 | 0 | BL27707 | 0.0166 | 0.0332 |
| BL06284 | 0.763 | 0 | BL01135 | 0.0166 | 0.0332 |

**Table 2.**
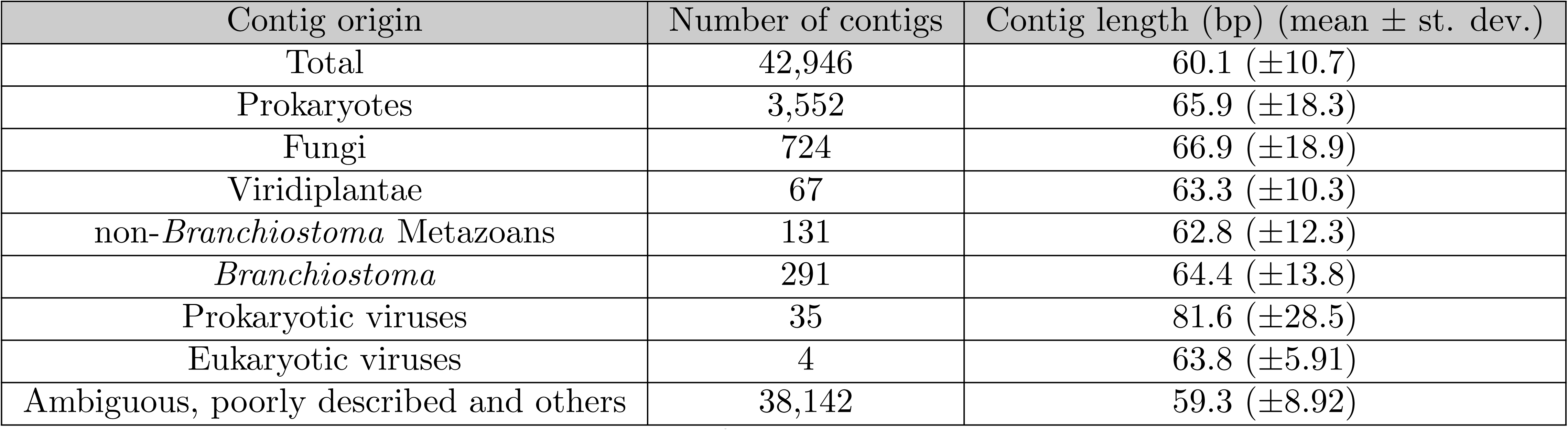
Annotation of reconstructed genomic contigs assembled from extragenomic, extratranscriptomic Small RNA-Seq reads.

It is formally possible that the few antisense exon-exon junction reads that we detected derive from an RNA polymerized by an RdRP. But their scarcity, as well as their extreme sequence homogeneity, suggests that they rather come from other sources (*e.g.*, DNA-dependent RNA polymerization, either from a *Branchiostoma* genomic locus or from a non-*Branchiostoma* contaminant) and map fortuitously on the BL05604 or BL00515 spliced transcript sequences. We note that *C. elegans* secondary siRNAs are highly diverse in sequence, and even low-throughput sequencing identifies antisense reads mapping on distinct exon-exon junctions [20]. We thus tend to attribute our observation of rare antisense exon-exon junction small RNAs to rare contaminants or sequencing errors, rather than to genuine RNA-dependent RNA polymerization in *Branchiostoma*.

Genes were sorted according to the total number of small RNA reads mapping on their exon-exon junctions in our pooled 24 Small RNA-Seq libraries. Numbers of mapped reads were normalized by the total number of genome-matching reads that do not match abundant non-coding RNAs, and expressed as *parts per million* (ppm). Top part: statistics for the whole transcriptome. Bottom part: only the top 10 genes in terms of sense-matching junction reads (left half of the table) or antisense-matching junction reads (right half of the table) are shown.

### Candidate *Branchiostoma* pathogens do not appear to be targeted by RNAi

In various other organisms, RNAi participates in the defence against pathogens (reviewed in [91]). Pathogen-specific siRNAs may exist in *Branchiostoma*, and they may have been too poorly abundant to be detected in our analyses of extragenomic, extratranscriptomic reads (see Supplementary File S1, section 5). We thus decided to interrogate specifically the populations of small RNAs mapping on *Branchiostoma* pathogen genomes. Several pathogenic bacteria (*Staphylococcus aureus, Vibrio alginolyticus* and *Vibrio anguillarum*; [92, 93]) have been described in various *Branchiostoma* species. We asked whether RNAi could target those pathogens *in vivo*. Focusing on the small RNA reads that do not map on the *Branchiostoma* genome or transcriptome, we observed large numbers of small RNAs deriving from these three bacterial genomes, indicating that the analyzed *Branchiostoma* specimens were in contact with those pathogens (after excluding reads that map simultaneously on 2 or 3 of these bacterial genomes, we detected 1,457,122 *S. aureus*-specific reads, 113,398 *V. alginolyticus*-specific reads and 103,153 *V. anguillarum*-specific reads in the pooled 24 Small RNA-Seq libraries; for reference: there are 125,550,314 *Branchiostoma* genome-matching reads in the pooled libraries). Small RNAs mapping on these pathogenic bacterial genomes do not display any obvious size distribution or sequence bias, thus suggesting that they constitute degradation products from longer bacterial RNAs rather than siRNAs (see Supplementary File S1, sections 7–9).

Our analyzed *Branchiostoma* specimens may also have been challenged by yet-unknown pathogens. Pooling every read that does not map on the *Branchiostoma* genome or transcriptome, across all 24 Small RNA-Seq libraries, offers the opportunity to reconstruct genomic contigs for the most abundant non-*Branchiostoma* sequences. In total, we collected 23,557,012 such extragenomic, extratranscriptomic reads. 42,946 contigs at least 50 bp long could be assembled from these reads using **velvet** [45]. Of these, 4,804 contigs could be annotated by homology search (see Table 2): 291 appear to match the *Branchiostoma* genome, and the reads supporting these contigs had probably failed to map properly on the genome because of sequencing errors or sequence polymorphism.

After assembling extragenomic, extratranscriptomic reads from all 24 libraries, contigs longer than 50 bp were annotated by a **blast** search on the NCBI non-redundant nucleic acid database.

We screened these contigs for potential *Branchiostoma* pathogens, which could be targeted by RNAi. Detected prokaryotic, fungal or non-*Branchiostoma* metazoan sequences may derive from symbiotic or commensal species rather than actual pathogens. Our analyzed adult specimens were collected from the natural environment, where unrelated organisms are expected to contaminate the samples; and our analyzed embryos were produced from gametes collected in non-sterile sea water. Following spawning, these gametes transit through the “atrium” (an open body cavity that putatively hosts various micro-organisms): so *in vitro*-fertilized embryos are also likely to be contaminated with non-pathogenic non-*Branchiostoma* species.

But we also observed several viral contigs, including 4 contigs from eukaryotic viruses. Three of them are matched by low numbers of small RNA reads, but the last one (a contig matching the *Acanthocystis turfacea* Chlorella virus 1 genome) is covered with high read counts in various developmental stages (see Supplementary Fig. S5). That virus is known to infect endosymbiotic algae of the protist *Acanthocystis turfacea*, and some reports suggest that it may also infect mammalian hosts [94], suggesting a broad tropism. Though still disputed [95, 96], this observation could suggest that *Branchiostoma* may also be sensitive to that virus. Yet, for this potential pathogen too, detected small RNA reads fail to display any size or sequence bias: they do not appear to be siRNAs (see Supplementary File S1, section 10).

Finally, we considered the possibility that some of the 38,142 un-annotated extragenomic contigs (see Table 2) may originate from unknown pathogens. We selected the 5 contigs displaying the highest read coverage (more than 200 ppm after pooling all 24 Small RNA-Seq libraries): small RNAs mapping on these hypothetical unknown pathogens also do not exhibit particular size or sequence biases, arguing against their nvolvement in RNAi (see Supplementary File S1, sections 11–15).

Because unambiguous RdRP-derived small RNAs could not be detected with certainty despite our efforts, and because we did not observe any small RNA population with classical siRNA size or sequence bias, we conclude that *Branchiostoma* RdRP genes are not involved in RNAi.

## Discussion

In cellular organisms, the only known function for RdRPs is the generation of siRNAs or siRNA precursors. It is thus frequently assumed [32, 47] or hypothesized [34] that animal RdRPs participate in RNAi. In particular, it has recently been proposed that arthropod RdRPs are required for RNAi amplification, and arthropod species devoid of RdRPs may rather generate siRNA precursors through bidirectional transcription [47]. While this hypothesis would provide an elegant explanation to the sporadicity of RdRP gene distribution in the phylogenetic tree, the provided evidence remains disputable: it has been proposed that a high ratio of antisense over sense RNA is diagnostic of bidirectional transcription, yet it remains to be explained why RNA-dependent RNA polymerization would produce less steady-state antisense RNA than DNA-dependent polymerization.

*Branchiostoma* 5’ monophosphorylated small RNAs do not appear to bear a 2’-*O* -methyl on their 3’ end: Libraries #2 contain few genome-matching sequences, and their size distribution suggests they are mostly constituted of contaminating RNA fragments rather than miRNAs, piRNAs or siRNAs. In every animal model studied so far, piRNAs were shown to bear a methylated 3’ end [25, 56–58, 85, 87, 97–99]. The enzyme responsible for piRNA methylation, Hen1 (also known as Pimet in *Drosophila*, HENN-1 in nematodes), has been identified in *Drosophila*, mouse, zebrafish and nematodes [55–58, 100–102]. In order to determine whether the absence of piRNA methylation in *Branchiostoma* could be due to an absence of the Hen1 enzyme, we searched for Hen1 orthologs in the predicted *Branchiostoma* proteome. Our HMMer search identified a candidate, BL03504. Its putative methyl-transferase domain contains every known important amino acid for Hen1 activity according to [103] (see Supplementary Fig. S6), suggesting that it is functional. Further studies will be required to investigate the biological activity of that putative enzyme, and to understand why it does not methylate *Branchiostoma* piRNAs.

Focusing on small RNA reads mapping on exon-exon junctions in the antisense orientation, we did not observe convincing evidence of RdRP activity in *Branchiostoma*. Even if RdRPs do not participate in RNAi, it could have been anticipated that Small RNA-Seq libraries could capture short degradation products of RdRP-polymerized long RNAs. This observation raises the possibility that the *Branchiostoma* RdRP genes do not express any active RdRP. At least these genes are transcribed: analysis of gene expression in long RNA-Seq data [43] shows a dynamic regulation, especially for the three genes with an intact predicted active site (see Supplementary Fig. S4).

One could hypothesize that these RdRPs do not play any biological function. Yet at least two of them, BL02069 and BL23385, possess a full-length RdRP domain with a preserved catalytic site. The conservation of these two intact genes suggests that they are functionally important. It can therefore be speculated that *Branchiostoma* RdRPs play a biological role, which is unrelated to RNAi. Such a function may involve the generation of double-stranded RNA (formed by the hybridization of template RNA with the RdRP product), but it could also involve single-stranded RdRP products. Future work will be needed to identify the biological functionality of these enzymes. We also note that the fungus *Aspergillus nidulans*, whose genome encodes two RdRPs with a conserved active site, does not require any of those for RNAi [104].

Animal RdRPs thus constitute an evolutionary enigma: not only have they been frequently lost independently in numerous animal lineages, but even in the clades where they have been conserved, their biological function seems to be variable. While RNAi is an ancient gene regulation pathway [1], involving the deeply conserved Argonaute and Dicer protein families, the role of RdRPs in RNAi appears to be accessory. Even though RdRPs are strictly required for RNAi in very diverse extant clades (ranging from nematodes to plants), it would be misleading to assume that RNAi constitutes their only biological function.

## Supporting information

**Fig. S1 Exclusion of dubious proteomes still indicates many independent RdRP losses.** (file ’Supplementary Figure after proteome selection.pdf’) Among the 538 analyzed proteomes, 442 contain at least 1,000 proteins of at least 1,000 amino acids (left panel) and 383 contain at least 5,000 proteins of at least 500 amino acids (right panel). Selective analysis of these species does not fundamentally change the results shown in Figure 1A. Same conventions than in Figure 1A. Some clades analyzed in Figure 1A could not be analyzed here after proteome exclusion: they are shown in grey.

**Fig. S2 Size and quality of the Small RNA-Seq libraries.** (file ’Supplementary Figure mapping statistics.pdf’) “No adapter” indicates hat the 3’ adapter was not detected in the read. “Extragenomic” means that the adapter-trimmed read does not match the *B. lanceolatum* genome assembly. “Abundant ncRNA” means that it maps on the genome assembly, on one of the genes for known abundant non-coding RNAs (rRNAs, tRNAs, snRNAs, snoRNAs, scaRNAs). “Genome mapper, not matching abundant ncRNAs” means that it maps elsewhere in the genome assembly.

**Fig. S3 Small RNA coverage in 15 hpf embryos for the two genes with highest antisense exon-exon junction read coverage.** (file ’Supplementary RdRP template loci with coverage.pdf’) Exons are represented by black rectangles. Detected small RNAs mapping on these genes in the sense orientation are shown in blue, those mapping in antisense orientation are in red. For antisense reads mapping on exon-exon junctions, their precise sequence (in red) is aligned with the gene sequence (in black; splicing donor and acceptor sites are in green).

**Fig. S4 Transcriptomics-based expression analysis of the 6 *Branchiostoma* RdRP genes.** (file ’Supplementary Figure S4.pdf’) For each of the six RdRP genes, mRNA abundance in various developmental stages was measured by RNA-Seq, and reported as cRPKM (corrected-for-mappability reads per kb and per million mapped reads; [105]). RdRP genes where an intact active site is predicted (see Fig. 1B) are annotated “with active site”. Adult RNA-Seq data is from NCBI’s BioSample accession #SAMN09381006 and SAMN09381007, other stages are from [43]. Adult male and female data were averaged. Temporal regulation of RdRP expression in embryos and juveniles was assessed by the Kruskal-Wallis test (*p*-values are indicated in the legend for each RdRP).

**Fig. S5 Small RNA coverage of the *Acanthocystis turfacea* Chlorella virus 1 (ATCV1) genome.** (file ’Supplementary Figure ATCV1 coverage.pdf’) *x* axis: genomic coordinate along the ATCV1 genome. *y* axis: number of reads covering each bp in the viral genome. Numbers of reads are expressed as *parts per million* (ppm) after normalization to the total number of *Branchiostoma* genome-matching reads that do not match abundant non-coding RNAs.

**Fig. S6 A *Branchiostoma* Hen1 candidate contains the known essential amino acids for Hen1 activity.** (file ’Supplementary Figure Hen1.pdf’) Sequences of 5 known Hen1 proteins (from *Nematostella vectensis, Danio rerio, Mus musculus, Arabidopsis thaliana* and *Drosophila melanogaster)* were aligned with the identified *Branchiostoma lanceolatum* Hen1 candidate (only the part of the alignment spanning 564 amino acids 661–939 of the *Arabidopsis* protein is shown). Alignment was performed 565 with **t-coffee** (version 11.00.8cbe486); other alignment programs (**Clustal Omega v.1.2.4**, **t-coffee v.8.93**, **Kalign v.2.03**, **MAFFT v.7.215**, but not **muscle v.3.8.31**) give the same main result: amino acids and amino acid combinations required for Hen1 catalytic activity [103] are conserved in the *Branchiostoma* candidate. Amino acids boxed in red were shown to be essential for *Arabodipsis* Hen1 activity; in orange: amino acids whose absence affects Hen1 activity without abolishing it entirely. Amino acid numbering is based on the *Arabidopsis* sequence.

**File S1. Size distribution and logo analyses of various small RNA classes.** (file ’Supplementary Data.pdf’) For each of the following classes, small RNA populations were analyzed as in Figure 3B, 3C and 4: reads matching the *B. lanceolatum* genome without matching abundant non-coding RNAs (section 1); reads matching *B. lanceolatum* pre-miRNA hairpins (section 2); reads matching the *B. lanceolatum* transcriptome without matching pre-miRNAs or abundant non-coding RNAs (section 3); reads matching *B. lanceolatum* mRNAs with long ORFs (section 4); reads not matching the *B. lanceolatum* genome or transcriptome (section 5); *C. elegans* small RNAs cloned with a procedure detecting 5’ mono- and polyphosphorylated RNAs [22] (section 6); reads not matching the *B. lanceolatum* genome or transcriptome, and matching the *Staphylococcus aureus* genome (section 7); reads not matching the *B. lanceolatum* genome or transcriptome, and matching the *Vibrio alginolyticus* genome (section 8); reads not matching the *B. lanceolatum* genome or transcriptome, and matching the *Vibrio anguillarum* genome (section 9); reads not matching the *B. lanceolatum* genome or transcriptome, and matching the *Acanthocystis turfacea* Chlorella virus 1 (ATCV1) genome (section 10); reads not matching the *B. lanceolatum* genome or transcriptome, and matching non-*Branchiostoma* contig #18690 (covered with 1,982.33 ppm small RNA reads across all 24 libraries) (section 11); reads not matching the *B. lanceolatum* genome or transcriptome, and matching non-*Branchiostoma* contig #7601 (covered with 1,534.35 ppm small RNA reads across all 24 libraries) (section 12); reads not matching the *B. lanceolatum* genome or transcriptome, and matching non-*Branchiostoma* contig #38312 (covered with 236.037 ppm small RNA reads across all 24 libraries) (section 13); reads not matching the *B. lanceolatum* genome or transcriptome, and matching non-*Branchiostoma* contig #3365 (covered with 223.535 ppm small RNA reads across all 24 libraries) (section 14); reads not matching the *B. lanceolatum* genome or transcriptome, and matching non-*Branchiostoma* contig #10883 (covered with 205.859 ppm small RNA reads across all 24 libraries) (section 15).

**Table S1 Detection of conserved miRNAs.** (file ’Supplementary Table 1.pdf’) *Branchiostoma lanceolatum* orthologs for *B. floridae* or *B. belcheri* pre-miRNA hairpins (as described in miRBase v.22) were screened for their predicted secondary structure and the abundance of the small RNAs they generate. Only those hairpins that comply with these rules are shown in this table. First column: name of orthologous pre-miRNA, and genomic coordinates in *B. lanceolatum*. Second column: sequences of the major forms of the 5’ arm and 3’ arm miRNAs, if expressed at ≥ 10 ppm in at least one developmental stage (miRNAs that do not meet that criterion are flagged “low abundance”). Third column: abundance of the 5’ arm and 3’ arm miRNAs in Libraries #1 along development. Embryonic stages contain mixed sexes; adult stages are shown in blue and pink for males and females, respectively. Trimming (up to 3 nt) and templated extension of miRNA 3’ ends were considered when measuring read counts.

## Supporting information

## Acknowledgments

The authors are grateful to Dr. Darryl Conte for helpful discussions and to Julie Claycomb, Kazufumi Mochizuki and Phillip D. Zamore for critical reading of the manuscript. We thank the *B. lanceolatum* genome consortium for the assembly and annotation of the *B. lanceolatum* genome, Dr. Manuel Irimia for assistance in transcriptomics analyses, and Dr. Ferdinand Marl’etaz for sharing unpublished data. This research was supported by an ATIP-Avenir grant from CNRS and Sanofi (to H.S.) and a post-doctoral fellowship from La Ligue contre le cancer (to N.P.). H.E.’s laboratory was supported by the CNRS and the ANR16-CE12-0008-01 and S.B. by the Institut Universitaire de France. We thank the MGX sequencing facility (Biocampus Montpellier, CNRS, INSERM, Univ. Montpellier, Montpellier, France). MGX acknowledges financial support from France G’enomique National infrastructure, funded as part of the “Investissement d’avenir” program managed by Agence Nationale pour la Recherche (contract ANR-10-INBS-09).

