## Supplementary figures and images for "Functional lability of RNA-dependent RNA polymerases in animals"

### Supplementary file 1

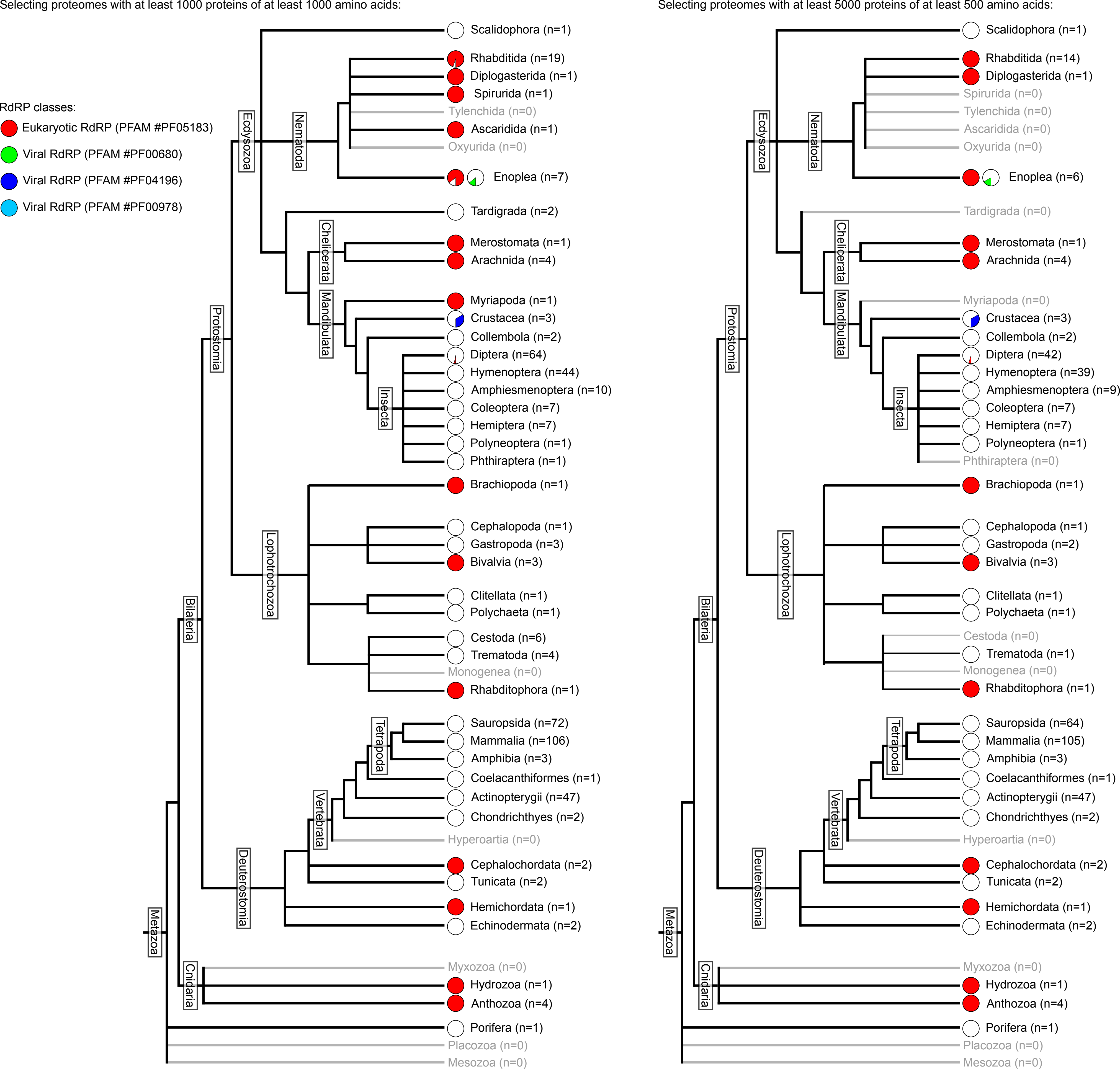

### Supplementary file 2

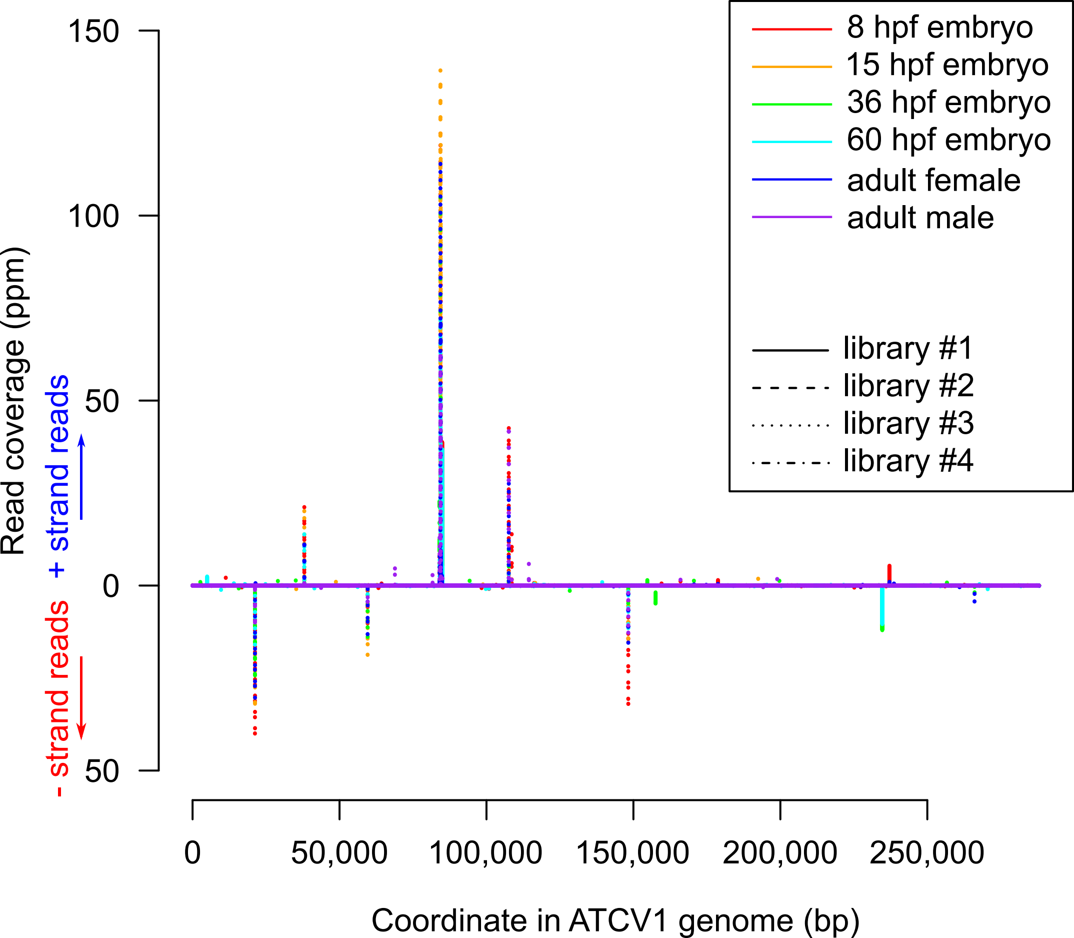

### Supplementary file 4

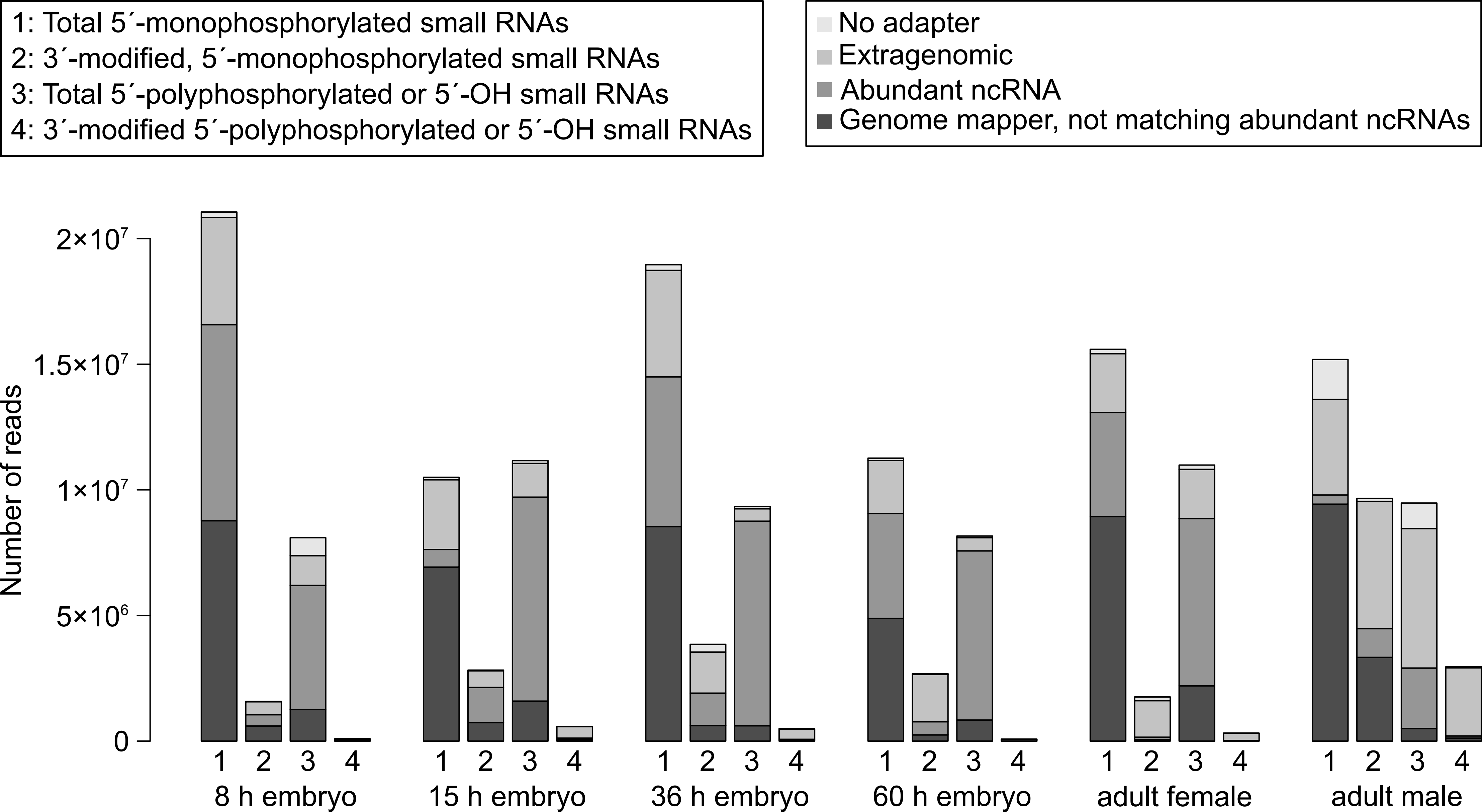

### Supplementary file 5

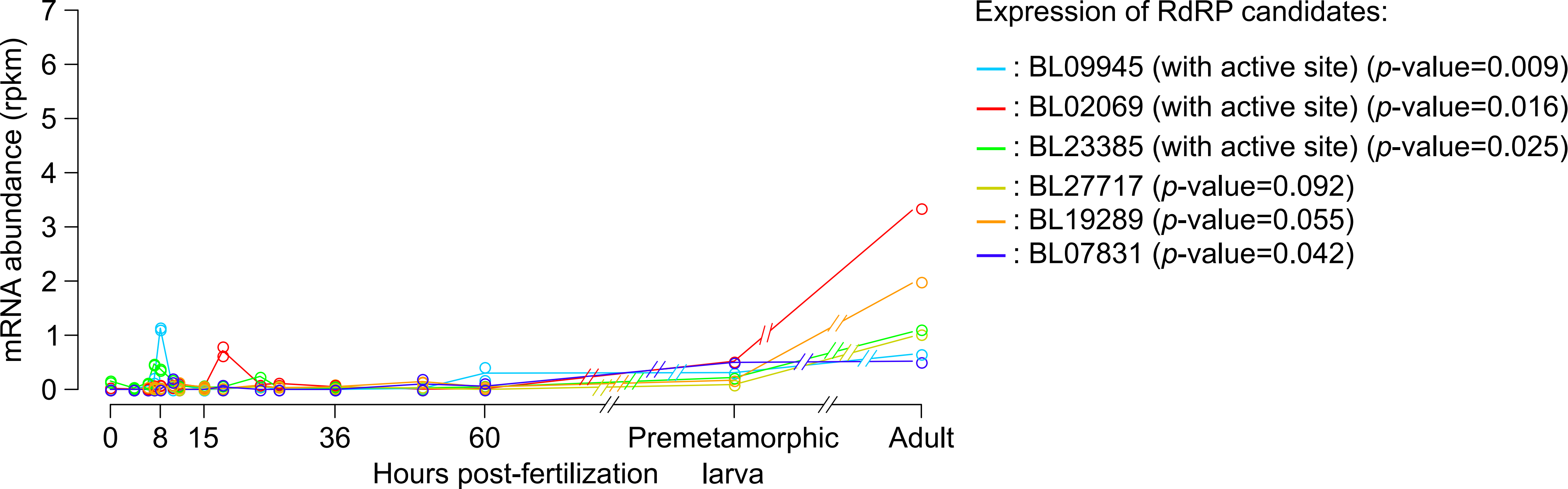

### Supplementary file 6

In 15 hpf embryos:

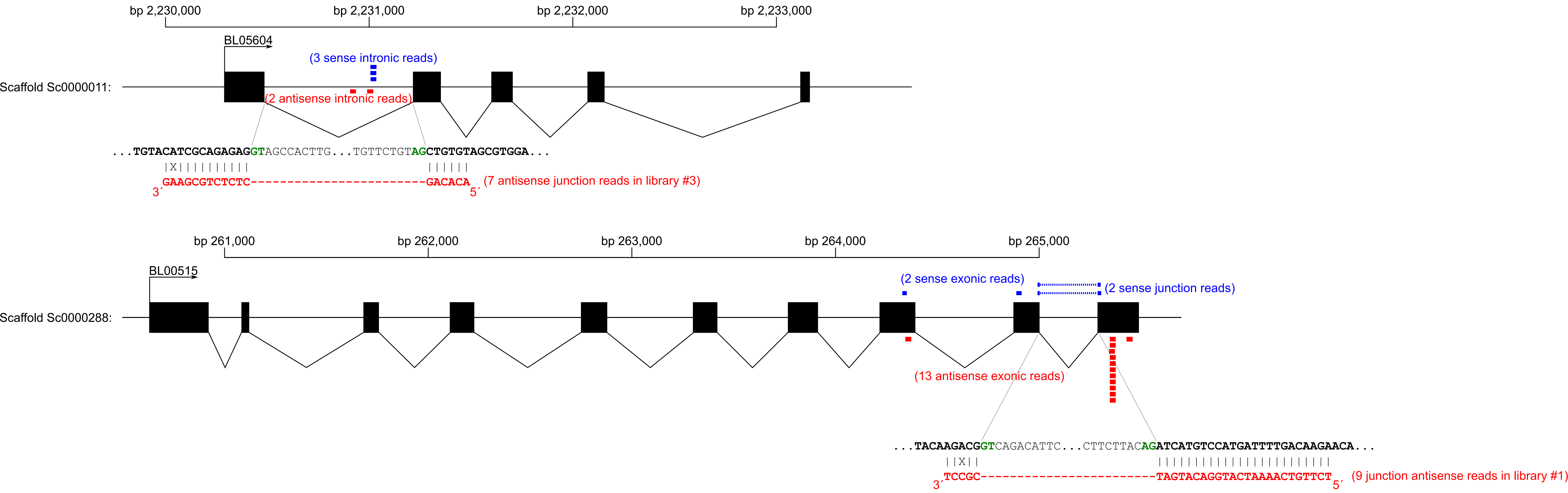
