## Supplementary material for "Functional lability of RNA-dependent RNA polymerases in animals"

```
                                                                 R701
B. lanceolatum BL03504     D-----------------------EEEFGGPVFSPPLYRQRYQTVADLVK----KYRPKR
N. vectensis AGW15602      -------------------------REQLGPKFDPPVYRQRYHRVIEVVK----EHKAKR
D. rerio NP_001017842      -----------------------------ATPFSPPLYMQRYQFVIDYVK----TYRPRK
M. musculus NP_001072114   E-----------------------VSPEKVIRFKPPLYKQRYQFVRDLVD----RHEPKK
A. thaliana NP_567616      IRSLLSERPCLNYNILLLGVKGPSEERMEAAFFKPPLSKQRVEYALKHIR----ESSAST
D. melanogaster NP_610732  ------------------------KMTETGITFDPPVYEQRYCATIQILEDARWKDQIKK

B. lanceolatum BL03504     LVDFGCAEGKLIRFLK-PEESLEQLTGIDLEGEVLESIRGIIKPLLSDYVQPRPRPFTVS
N. vectensis AGW15602      VLDFGCAEAKMLRSLINSTTNIEELVGVDIDRDLLEDSIFRIRPLTTDYLTPRPHPLAVS
D. rerio NP_001017842      VIDFGCAECCLLKKLKFHRNGIQLLVGVDINSVVLLKRMHSLAPLVSDYLQPSDGPLTIE
M. musculus NP_001072114   VADLGCGDAKLLKLLKI-YPCIQLLVGVDINEEKLHSNGHRLSPYLGEFVKPRDLDLTVT
A. thaliana NP_567616      LVDFGCGSGSLLDSLLDYPTSLQTIIGVDISPKGLARAAKMLHVKLN---KEACNVKSAT
D. melanogaster NP_610732  VVEFGCAEMRFFQLMR-RIETIEHIGLVDIDKSLLMRNLTSVNPLVSDYIRSRASPLKVQ

                                              E796     E799, H800
B. lanceolatum BL03504     LYQGSIAECDDRFKSYDMVTCVEVIEHLDPPVLDAMPSNVFGHMRPSVVVVTTPNSEFNV
N. vectensis AGW15602      LYQGSISKADDRFCDFDVVACIEIVEHLVPEHLEAMPAVLLGQLSPLVAIVTTPNADFNV
D. rerio NP_001017842      LYQGSVMEREPCTKGFDLVTCVELIEHLELEEVERFSEVVFGYMAPGAVIVTTPNAEFNP
M. musculus NP_001072114   LYHGSVVERDSRLLGFDLITCIELIEHLDSDDLARFPDVVFGYLSPAMVVISTPNAEFNP
A. thaliana NP_567616      LYDGSILEFDSRLHDVDIGTCLEVIEHMEEDQACEFGEKVLSLFHPKLLIVSTPNYEFNT
D. melanogaster NP_610732  ILQGNVADSSEELRDTDAVIAIELIEHVYDDVLAKIPVNIFGFMQPKLVVFSTPNSDFNV

                                                         H860
B. lanceolatum BL03504     LFPN--------------F-----SGFRNADHRFEWTRQEFQTWAEGVAQRF-SYDVTFH
N. vectensis AGW15602      LFPD--------------L-----VGFRHWDHKFEWTRAEFKDWATSQADKF-GYSVTFE
D. rerio NP_001017842      LLPG--------------L-----RGFRNYGHKFEWTRAEFQTWAHRVCREH-GYSVQFT
M. musculus NP_001072114   LFPT--------------V------TLRDADHKFEWSRMEFQTWALHVANCY-NYRVEFT
A. thaliana NP_567616      ILQRSTPETQEENNSEPQL-----PKFRNHDHKFEWTREQFNQWASKLGKRH-NYSVEFS
D. melanogaster NP_610732  IFTR--------------FNPLLPNGFRHEDHKFEWSRDEFKNWCLGIVEKYPNYMFSLT

B. lanceolatum BL03504     GIGTGPEGTEHLGCCTQMAIFERKQTPYDE--------N-STVLWGTPYELIAEAVFPYR
N. vectensis AGW15602      GIGSGPSGTEHLGCCSQMALFIKQNTAPA---------G-RQTGFGEPYNLIARVEHPYR
D. rerio NP_001017842      GVGEAAGHWRDVGFCTQIAVFQRNFDGVNRSMS-----N-AEHLEPSVYRLLYRVVYPSL
M. musculus NP_001072114   GVGTPPAGSEHVGYCTQIGVFTKNGGKLSK-PS-----V-SQQCDQHVYKPVYTTSYPSL
A. thaliana NP_567616      GVGGS--GEVEPGFASQIAIFRREASSVE---------N-VAESSMQPYKVIWEWKKEDV
D. melanogaster NP_610732  GVGNPPKEYESVGPVSQIAIFVRKDMLEMQ-LVNPLVSKPNIDKESIPYKLIHTVEYPFY

B. lanceolatum BL03504     ENTLSKEQQILQEVQYYIRQIM--QRRVHGEDEEKD---NADDDDSAPE-----------
N. vectensis AGW15602      KCTLTEEEKILIELDRTLWFLS--QPSA-YEDDEISD--SEDLGD--DK-----------
D. rerio NP_001017842      CDNNIYQKTLINEVLYEAQHLR--QQWL-IRENMNNN----AHFYS---P-PLMEALHHG
M. musculus NP_001072114   QQEKVLKFVLVGELLIQVDRLRLRYQRM-LRDREKDRGPKPGDMDSCPAPHLLLGAVFTE
A. thaliana NP_567616      ---------------------------------EKKK-----------------------
D. melanogaster NP_610732  VDTRTEKEKLWTEVQIELQRFK--RQF---ESSEIEE---GTYQDT--------------
```
