## Supplementary material for "Functional lability of RNA-dependent RNA polymerases in animals"

| Pre-miRNA | miRNA sequences | Abundance profile in development |
|---|---|---|
| bfl-mir-71 ortholog at Sc0000005 bp 2992213-2992115 (- strand) | 5´arm: `UGAAAGACAUGGGUAGUGAGAU`<br><br>3´arm: `CCCAUUUCCCUGUCUUUCAAC` | 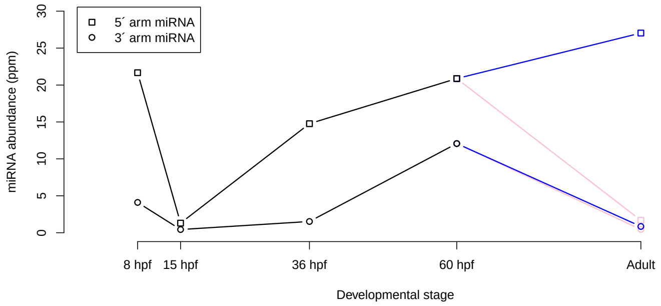 |
| bfl-mir-4856b ortholog at Sc0000022 bp 2102339-2102425 (+ strand) | 5´arm: `UCGCAUUGACGUCAGCGCCGUU`<br><br>3´arm: `(low abundance)` | 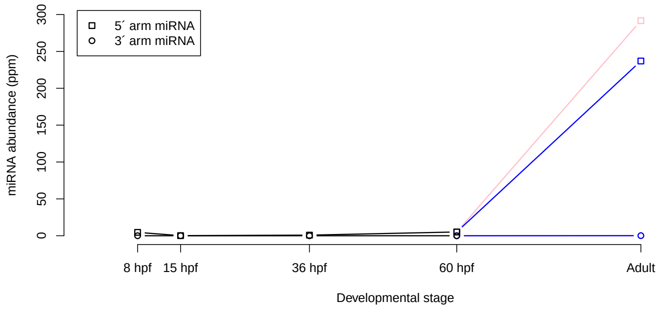 |
| bfl-mir-4876 ortholog at Sc0000005 bp 2219860-2219951 (+ strand) | 5´arm: `(low abundance)`<br><br>3´arm: `CUUACGUGCCACGUGAGGACUCU` | 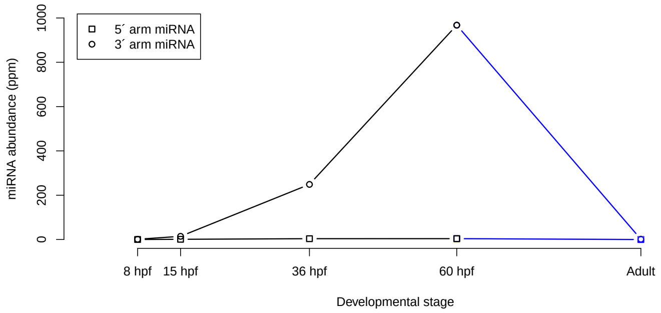 |
| bfl-mir-10c ortholog at Sc0000000 bp 2027699-2027621 (- strand) | 5´arm: `UACCCUGUAGAUCCGGACUUGUGA`<br><br>3´arm: `(low abundance)` | 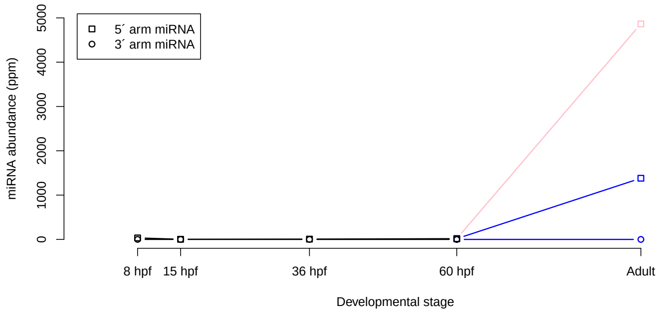 |

(continued on next page)

| Pre-miRNA | miRNA sequences | Abundance profile in development |
|---|---|---|
| bfl-mir-4869 ortholog at Sc0000022 bp 2099003-2099080 (+ strand) | 5´arm: `ACGAUGUUGACUUCCGCUCCUCU`<br>3´arm: `GGGACCUGUAGUCAACACGAGA` | 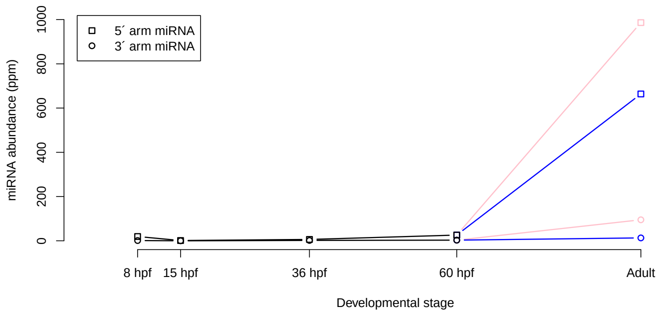 |
| bbe-mir-125a ortholog at Sc0000265 bp 98448-98548 (+ strand) | 5´arm: `UCCCUGAGACCCUAACUUGUGA`<br>3´arm: `ACAGGUUAGGAUCUUGGGAGCU` | 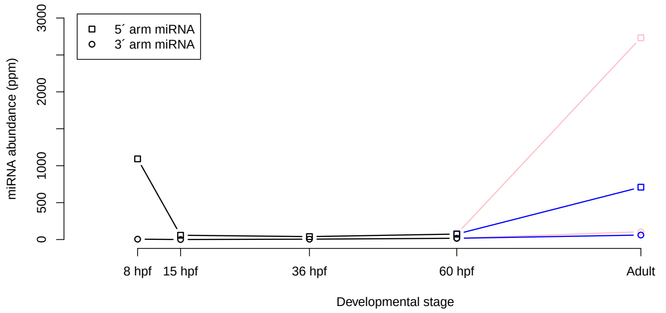 |
| bbe-mir-4874 ortholog at Sc0000043 bp 1147175-1147086 (- strand) | 5´arm: `AGUUUGUAGCAUCCAGCUGAGCU`<br>3´arm: `CUGGCUGGUUGGUGCAAACAGG` | 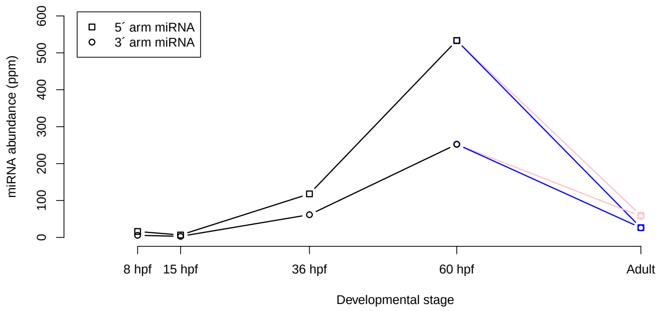 |
| bbe-mir-281 ortholog at Sc0000000 bp 9083742-9083658 (- strand) | 5´arm: `AGGAGAGCCGUUCUGUGACUGU`<br>3´arm: `(low abundance)` | 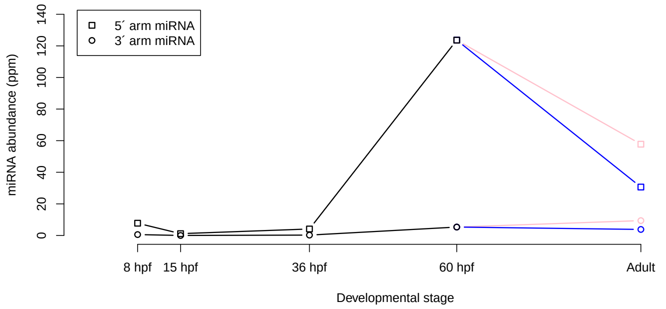 |

(continued on next page)

<table>
<tr><th>Pre-miRNA</th><th>miRNA sequences</th><th>Abundance profile in development</th></tr>
<tr><td>bbe-mir-92a-2 ortholog at Sc0000007 bp 4735603-4735697 (+ strand)</td><td>5´arm:<br>AGGCCAGGAUUGGUGGCAAUGCC<br><br>3´arm:<br>UAUUGCACUUGUCCCGGCCUUU</td><td>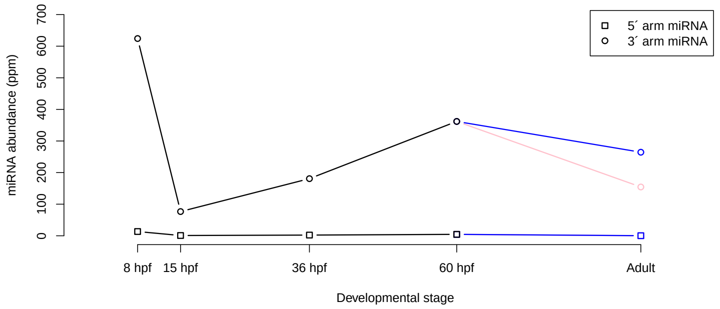
</td></tr>
<tr><td>bbe-mir-2056 ortholog at Sc0000086 bp 322130-322049 (- strand)</td><td>5´arm:<br>CAGGUAUGUCUGCGGUGAGGCU<br><br>3´arm:<br>UUUCACUGUAGAUCUACCUGCG</td><td>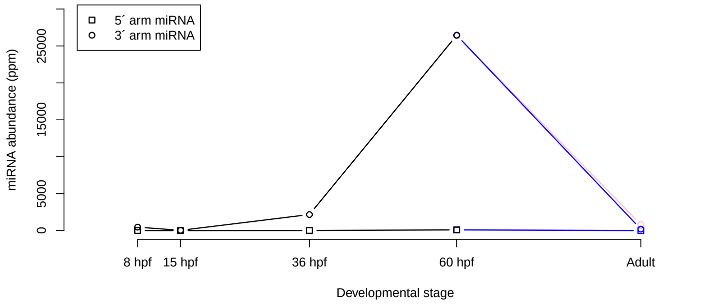
</td></tr>
<tr><td>bbe-mir-4880 ortholog at Sc0000057 bp 1180856-1180942 (+ strand)</td><td>5´arm:<br>UUUGCUAUUCGAUGACCAGUGG<br><br>3´arm:<br>(low abundance)</td><td>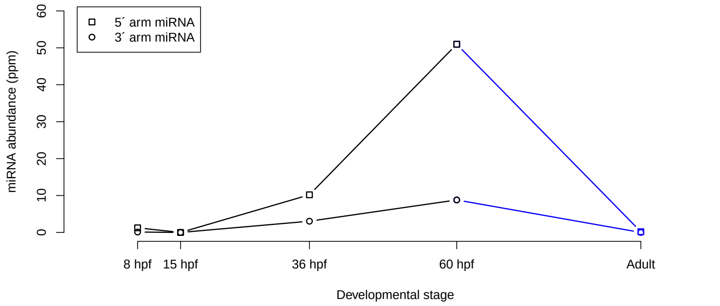
</td></tr>
<tr><td>bbe-mir-31 ortholog at Sc0000399 bp 174399-174498 (+ strand)</td><td>5´arm:<br>UGGCAAGAUGUUGGCAUAGCUG<br><br>3´arm:<br>(low abundance)</td><td>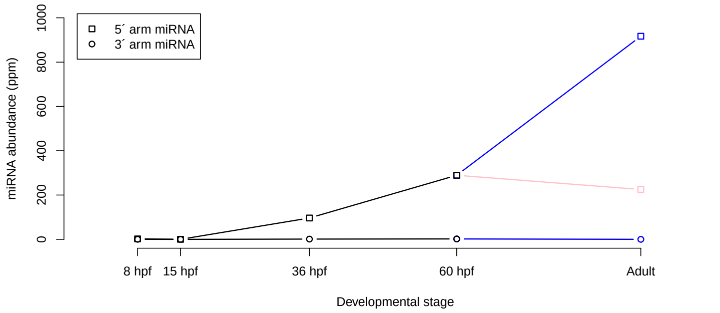
</td></tr>
</table>

(continued on next page)

| Pre-miRNA | miRNA sequences | Abundance profile in development |
|---|---|---|
| bfl-mir-200b ortholog at Sc0000010 bp 3877902-3877999 (+ strand) | 5´arm: `(low abundance)`<br><br>3´arm: `UAAUACUGUCUGGUAAUGAUGUU` | 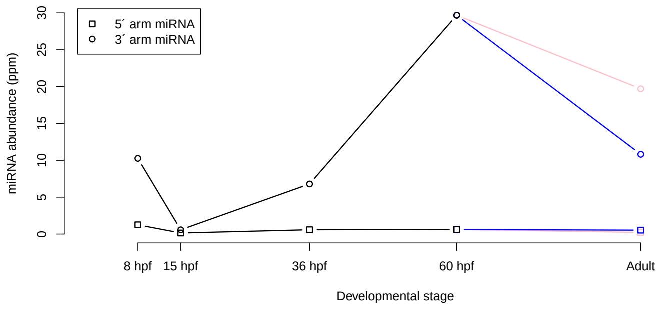 |
| bfl-mir-2071 ortholog at Sc0000288 bp 188813-188893 (+ strand) | 5´arm: `AUGCGGUGCGGUGGUAGCAACCG`<br><br>3´arm: `AUUGUUACACCGCGCCGCAAAAG` | 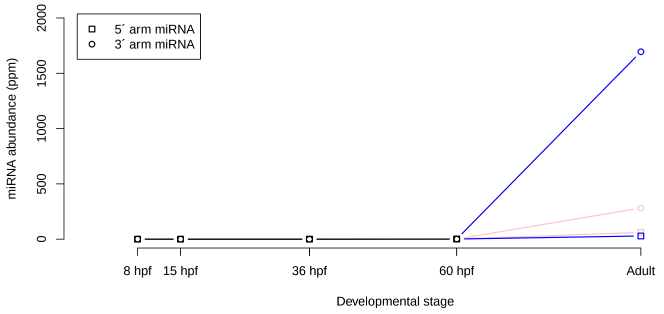 |
| bfl-mir-4865 ortholog at Sc0000063 bp 315717-315798 (+ strand) | 5´arm: `(low abundance)`<br><br>3´arm: `UGUAGAGAGAGUGACAGGUUGU` | 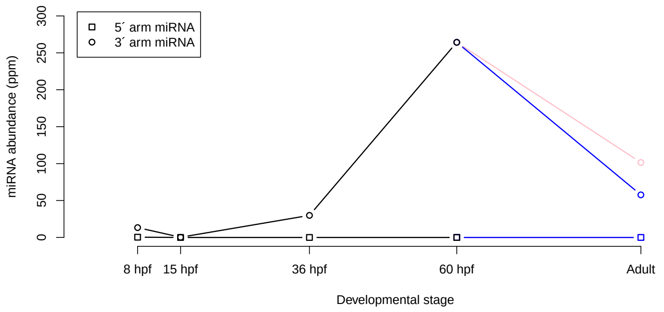 |
| bbe-mir-200c ortholog at Sc0000010 bp 3875272-3875359 (+ strand) | 5´arm: `(low abundance)`<br><br>3´arm: `UAACACUGUCUGGUAAUGAUG` | 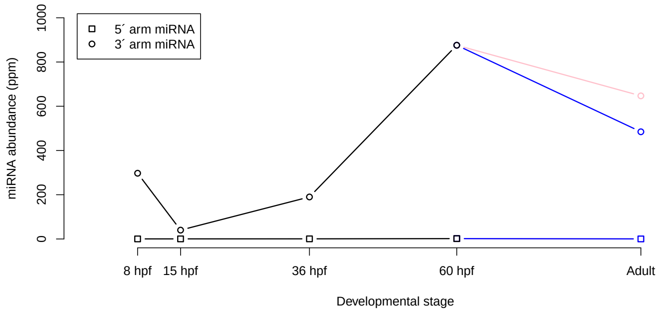 |

(continued on next page)

| Pre-miRNA | miRNA sequences | Abundance profile in development |
|---|---|---|
| bfl-mir-29a ortholog at Sc0000043 bp 1849481-1849575 (+ strand) | 5´arm: `GCUGAUUUCAGUUGGUGCUAGA`<br><br>3´arm: `(low abundance)` | 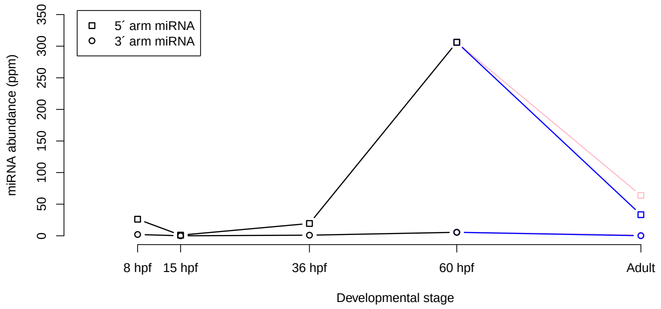 |
| bbe-mir-2058 ortholog at Sc0000110 bp 921627-921549 (- strand) | 5´arm: `UGAGAAGUAAGACUACCAUCCCGU`<br><br>3´arm: `(low abundance)` | 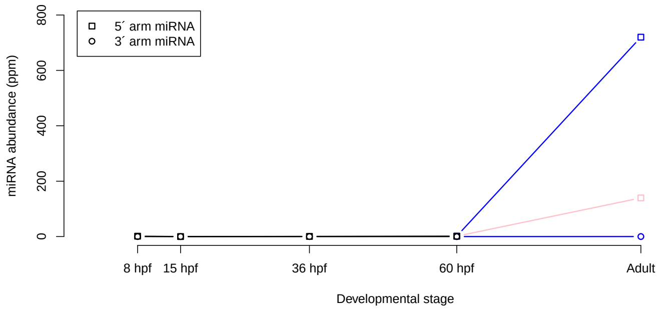 |
| bfl-let-7a-2 ortholog at Sc0000265 bp 98758-98852 (+ strand) | 5´arm: `UGAGGUAGUAGGUUGUAUAGUU`<br><br>3´arm: `CUGUGCAACCUGCUAGCUCUCC` | 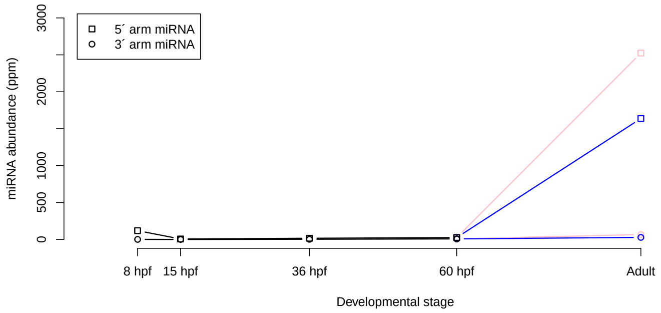 |
| bfl-mir-200c ortholog at Sc0000010 bp 3875267-3875358 (+ strand) | 5´arm: `(low abundance)`<br><br>3´arm: `UAACACUGUCUGGUAAUGAUG` | 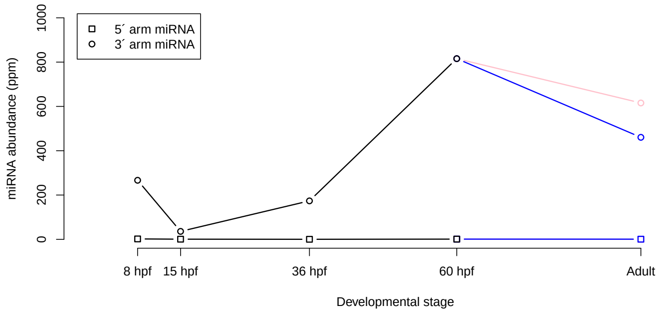 |

(continued on next page)

| Pre-miRNA | miRNA sequences | Abundance profile in development |
|---|---|---|
| bfl-mir-4860 ortholog at Sc0000063 bp 316331-316404 (+ strand) | 5´arm: UGCCUGUCAACGUCUCUGUACA<br><br>3´arm: UGUAGAGAUUGUGUGACGGGUAGU | 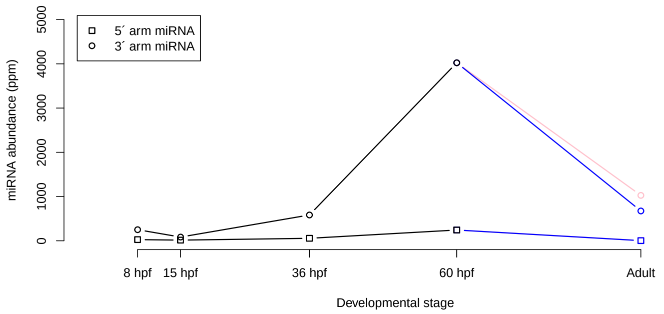 |
| bfl-mir-4864 ortholog at Sc0000184 bp 79034-79113 (+ strand) | 5´arm: AGGGAGAUCGUCUCGGGCAUACA<br><br>3´arm: UAGCCAGACCUGAUCUCCCUGC | 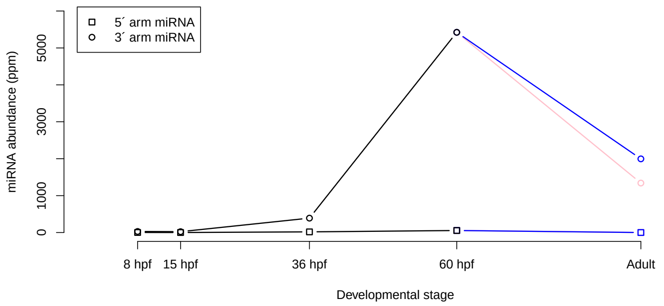 |
| bfl-mir-4861 ortholog at Sc0000005 bp 3994270-3994362 (+ strand) | 5´arm: AGCCAAUGCGGCAUGUAAAAGGC<br><br>3´arm: UUUACGUGCCACAUUGUCUCCU | 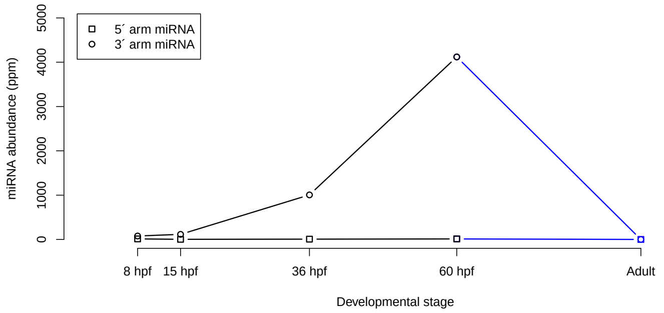 |
| bfl-mir-2062 ortholog at Sc0000099 bp 755622-755703 (+ strand) | 5´arm: UGCAACAAUAUUUCAUCAGUGG<br><br>3´arm: ACUGGUGAAAUGUAGUUGCGUA | 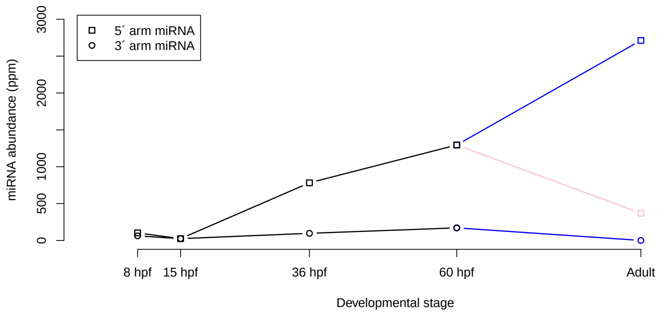 |

(continued on next page)

| Pre-miRNA | miRNA sequences | Abundance profile in development |
|---|---|---|
| bbe-mir-4859 ortholog at Sc0000221 bp 303532-303447 (- strand) | 5´arm: AGCAGCGAGCAUUACGGUCAUU<br>3´arm: UGACAGUAAUGCCCGCUGACUU | 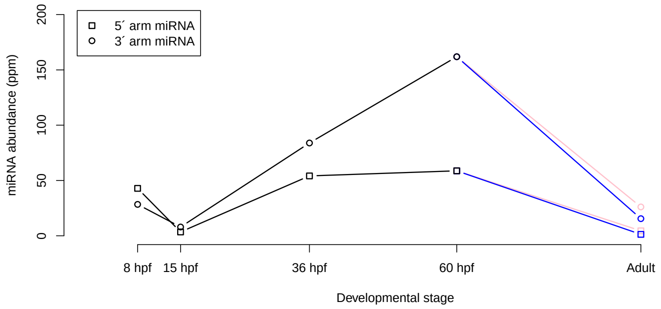 |
| bfl-mir-4871 ortholog at Sc0000001 bp 4527403-4527475 (+ strand) | 5´arm: UCUGAAGUACCUGUUGCCAAAGG<br>3´arm: UUUGGCACUGGUACUUUGGAGU | 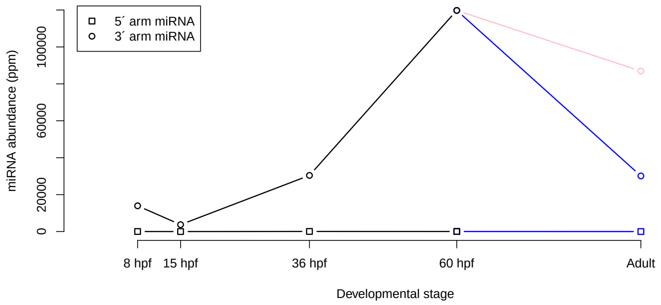 |
| bbe-mir-100 ortholog at Sc0000265 bp 96682-96781 (+ strand) | 5´arm: AACCCGUAGAUCCGAACUUGUGU<br>3´arm: CAAGCUCGUGUCUAUGGGUCU | 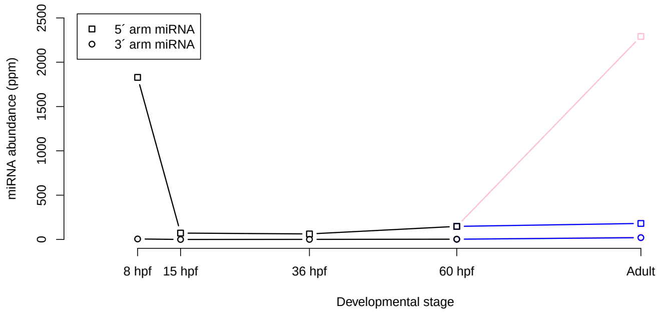 |
| bbe-mir-22 ortholog at Sc0000015 bp 676722-676797 (+ strand) | 5´arm: AGCUCUUCACUCGGUAGCUCUG<br>3´arm: AAGCUGCCAGAUGAAGAGCUGU | 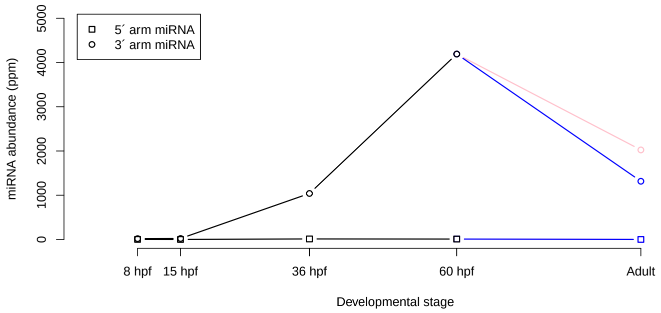 |

(continued on next page)

| Pre-miRNA | miRNA sequences | Abundance profile in development |
|---|---|---|
| bfl-mir-190 ortholog at Sc0000166 bp 638463-638558 (+ strand) | 5´arm: `UGAUAUGUUUGAUAUUUGGUUG`<br>3´arm: `(low abundance)` | 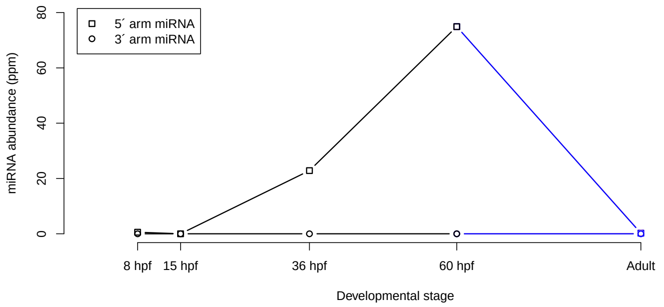 |
| bbe-mir-2057 ortholog at Sc0000110 bp 922600-922519 (- strand) | 5´arm: `UGAGAAGUUAGCCAACCAUCCGG`<br>3´arm: `(low abundance)` | 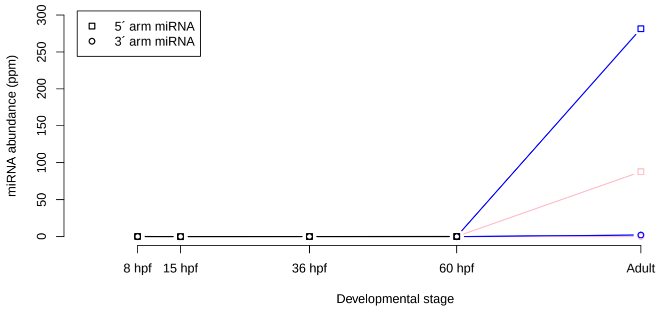 |
| bbe-mir-2061 ortholog at Sc0000005 bp 2547710-2547629 (- strand) | 5´arm: `(low abundance)`<br>3´arm: `UUGCAUAGGUACAUUGGUCAGU` | 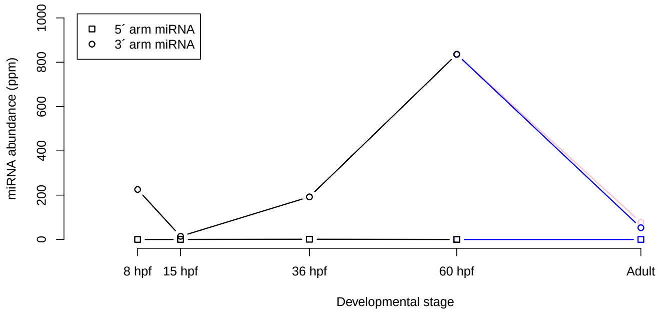 |
| bfl-mir-124 ortholog at Sc0000076 bp 963692-963792 (+ strand) | 5´arm: `AGUGUUCACGGCGGUCCUUAAU`<br>3´arm: `(low abundance)` | 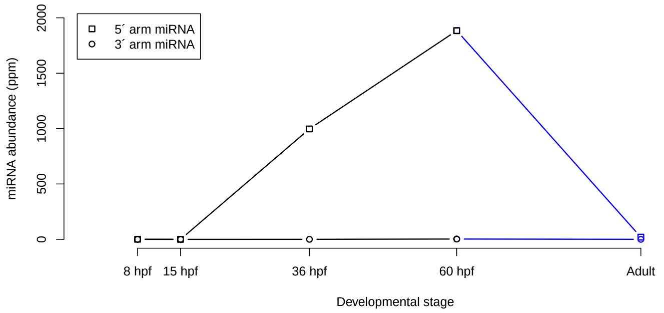 |

(continued on next page)

<table>
<tr><th>Pre-miRNA</th><th>miRNA sequences</th><th>Abundance profile in development</th></tr>
<tr><td>bfl-mir-4873 ortholog at Sc0000015 bp 1919187-1919110 (- strand)</td><td>5´arm: <code>UGUUCCCACCUUCUGAUGUUGUU</code><br><br>3´arm: <code>(low abundance)</code></td><td>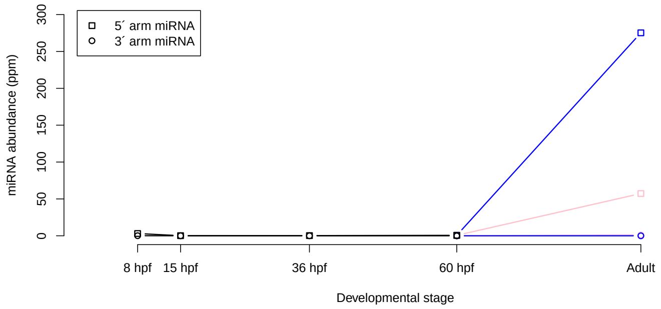
</td></tr>
<tr><td>bfl-mir-4891 ortholog at Sc0000002 bp 2469330-2469421 (+ strand)</td><td>5´arm: <code>(low abundance)</code><br><br>3´arm: <code>CGUACCAGGACGCUCGUCUGCC</code></td><td>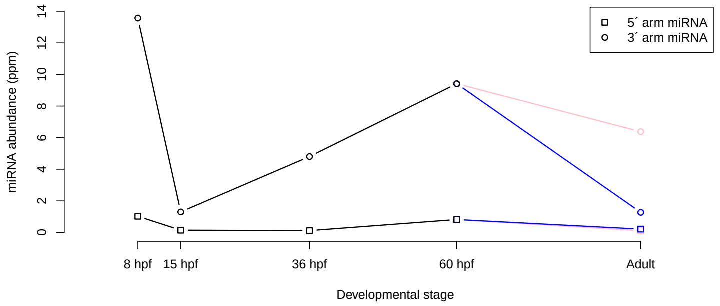
</td></tr>
<tr><td>bfl-mir-2070 ortholog at Sc0000079 bp 1008448-1008530 (+ strand)</td><td>5´arm: <code>UUUCCACAGCCUCUACACAUGU</code><br><br>3´arm: <code>AUGUGCAUAAGCUGUGGGAGCA</code></td><td>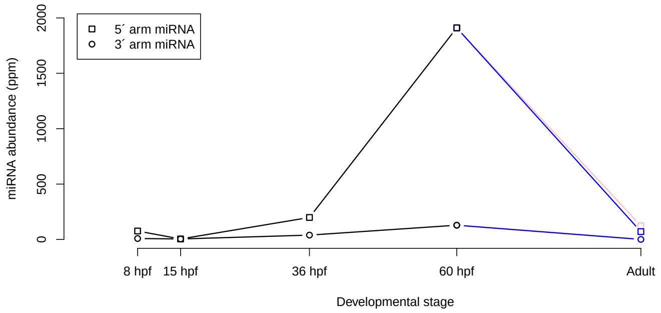
</td></tr>
<tr><td>bfl-mir-2076 ortholog at Sc0000004 bp 4642959-4643036 (+ strand)</td><td>5´arm: <code>AAUUGCACUAGAGUGAUUUGUU</code><br><br>3´arm: <code>(low abundance)</code></td><td>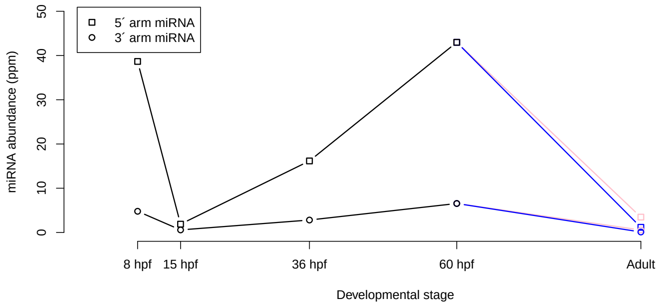
</td></tr>
<tr><td colspan="3" align="right">(continued on next page)</td></tr>
</table>

| Pre-miRNA | miRNA sequences | Abundance profile in development |
|---|---|---|
| bbe-mir-133 ortholog at Sc0000092 bp 690262-690162 (- strand) | 5´arm: `AAAGCUGGUAAAUUGGAACCA` 3´arm: `UUGGUCCCCUUCAACCAGCUGU` | 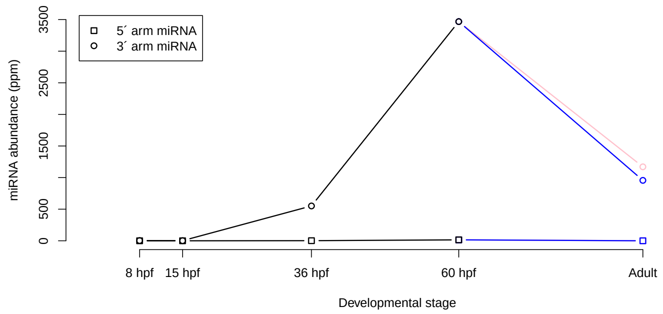 |
| bfl-mir-92b ortholog at Sc0000007 bp 2890460-2890560 (+ strand) | 5´arm: `AGGUCUGGACAGUUGCAAUCUU` 3´arm: `CAUUGCACUCGUCCCGGCCUGA` | 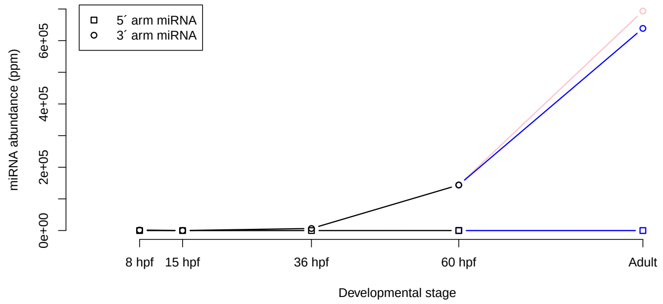 |
| bfl-mir-137 ortholog at xpSc0039671 bp 309878-309778 (- strand) | 5´arm: `(low abundance)` 3´arm: `UAUUGCUUGAGAAUACACGUGA` | 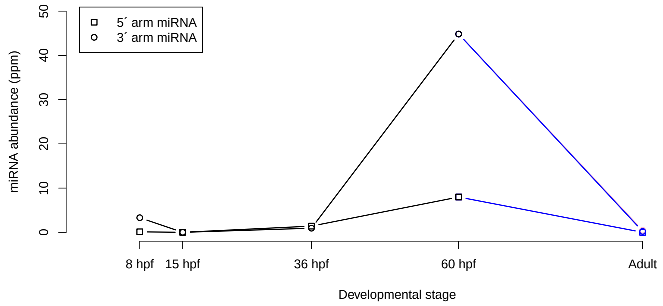 |
| bfl-mir-4868b ortholog at Sc0000017 bp 1346380-1346293 (- strand) | 5´arm: `CUCAUCACACCGGAAGCUGUUA` 3´arm: `UCAGCUCCAGCUGUGAUGAGUG` | 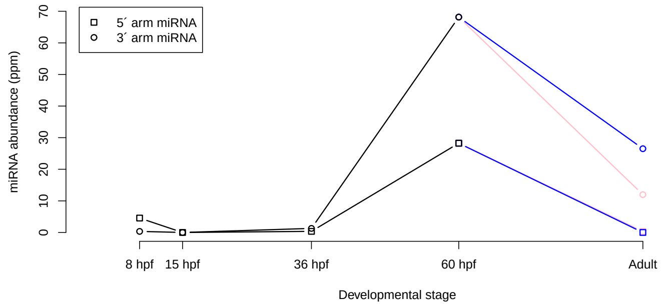 |

(continued on next page)

| Pre-miRNA | miRNA sequences | Abundance profile in development |
|---|---|---|
| bfl-mir-242 ortholog at Sc0000023 bp 500019-499937 (- strand) | 5´arm:<br>`UUGCGUAGGCGUUGUGCACACU`<br><br>3´arm:<br>`(low abundance)` | 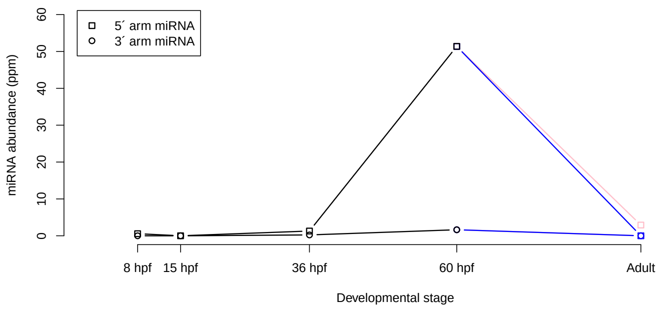 |
| bfl-mir-92d ortholog at Sc0000062 bp 529331-529413 (+ strand) | 5´arm:<br>`(low abundance)`<br><br>3´arm:<br>`UAUUGCACUUAUCCUGGCCUGU` | 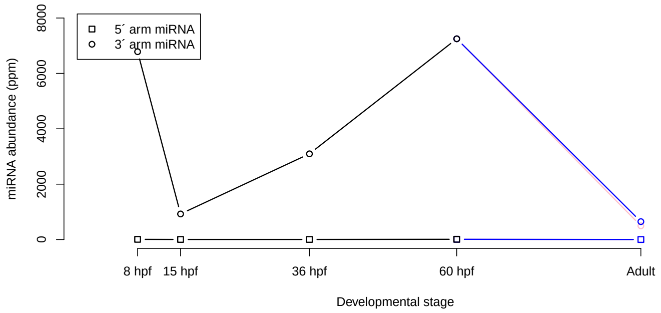 |
| bfl-mir-4880 ortholog at Sc0000057 bp 1180852-1180940 (+ strand) | 5´arm:<br>`UUUGCUAUUCGAUGACCAGUGG`<br><br>3´arm:<br>`(low abundance)` | 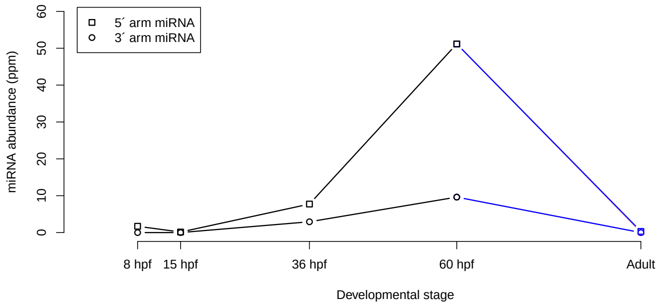 |
| bbe-mir-92a-1 ortholog at Sc0000007 bp 4735908-4735986 (+ strand) | 5´arm:<br>`AGGUCGUGAUAGGCGGCAAUGUU`<br><br>3´arm:<br>`UAUUGCACUUGUCCCGGCCUUU` | 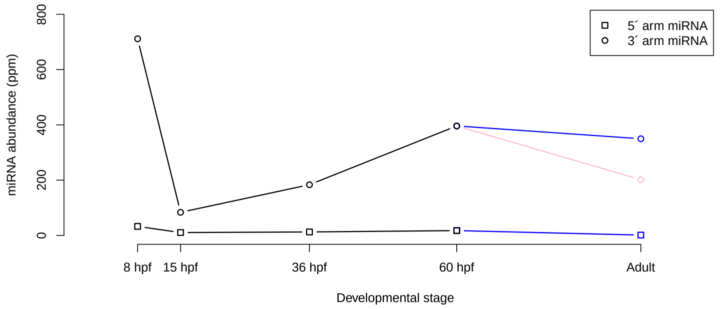 |

(continued on next page)

| Pre-miRNA | miRNA sequences | Abundance profile in development |
|---|---|---|
| bfl-mir-2058 ortholog at Sc0000110 bp 921624-921546 (- strand) | 5´arm: `UGAGAAGUAAGACUACCAUCCCGU`<br>3´arm: `(low abundance)` | 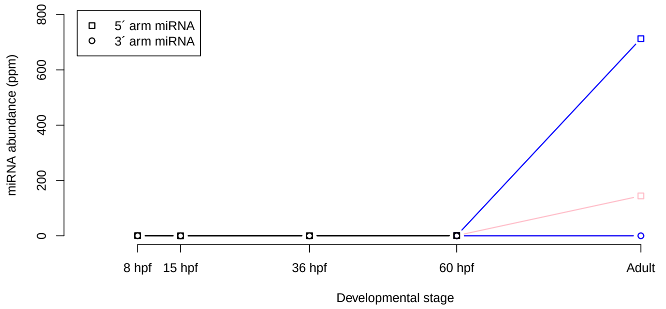 |
| bfl-mir-10b ortholog at Sc0000000 bp 1938354-1938254 (- strand) | 5´arm: `AACCCUGUGGAUCCGAUCUUGUG`<br>3´arm: `(low abundance)` | 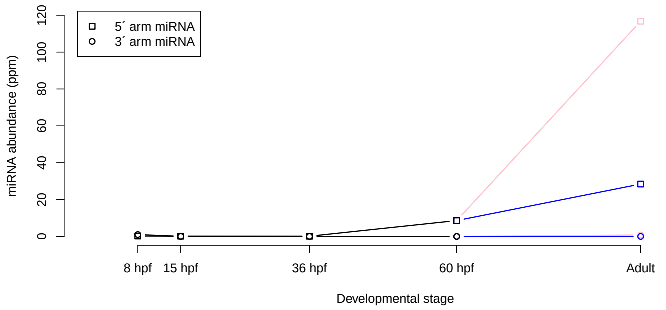 |
| bfl-mir-4856a ortholog at Sc0000022 bp 2103002-2103085 (+ strand) | 5´arm: `ACGCAGUGACGUCAGCGCCUCU`<br>3´arm: `(low abundance)` | 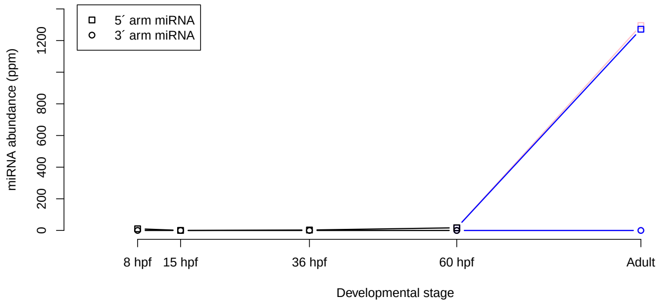 |
| bfl-mir-183 ortholog at Sc0000043 bp 398562-398462 (- strand) | 5´arm: `UAUGGCACUGGUAGAAUUCACUGA`<br>3´arm: `(low abundance)` | 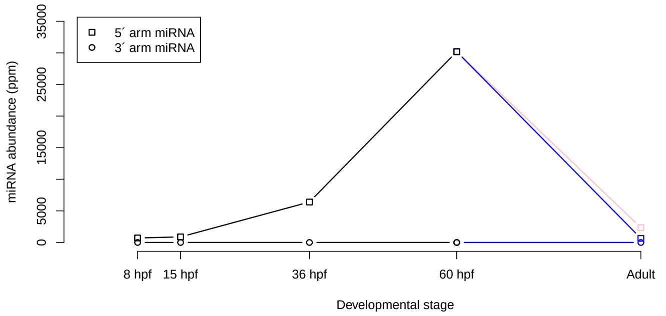 |

(continued on next page)

| Pre-miRNA | miRNA sequences | Abundance profile in development |
|---|---|---|
| bbe-mir-2067 ortholog at Sc0000082 bp 843448-843367 (- strand) | 5´arm:<br>`(low abundance)`<br><br>3´arm:<br>`AAGCAGCAGUAUGCAAUGGUGA` | 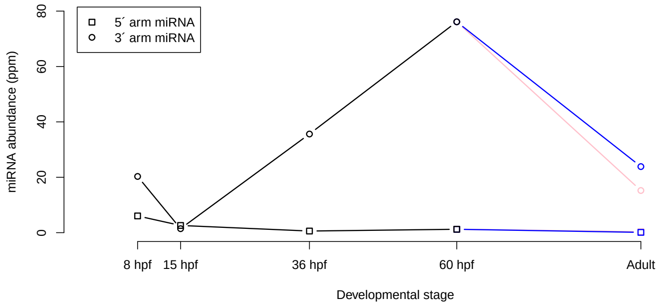 |
| bfl-mir-184 ortholog at Sc0000017 bp 1306111-1306211 (+ strand) | 5´arm:<br>`CUUAUCACUUCUUCCGCCGAGC`<br><br>3´arm:<br>`UGGACGGAGAACUGAUAAGGGCC` | 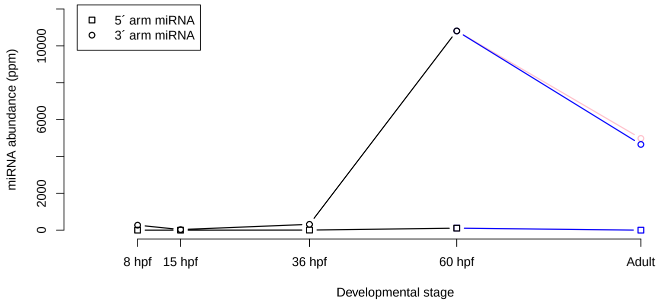 |
| bbe-mir-34a ortholog at Sc0000034 bp 210967-210880 (- strand) | 5´arm:<br>`CGUUCCCUGUGUGCUGCUG`<br><br>3´arm:<br>`AGCCACUGUACACACUCCCGUA` | 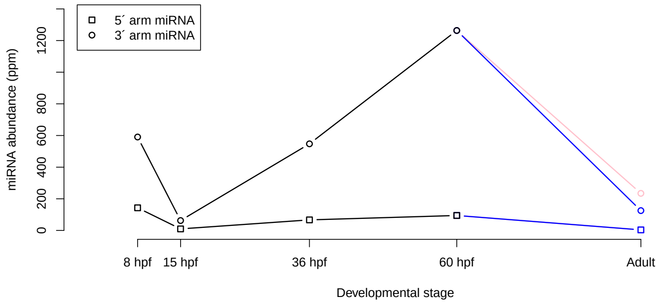 |
| bfl-mir-4904 ortholog at Sc0000028 bp 432156-432080 (- strand) | 5´arm:<br>`UCCCGGAGUCGUUCUAUACGCCG`<br><br>3´arm:<br>`(low abundance)` | 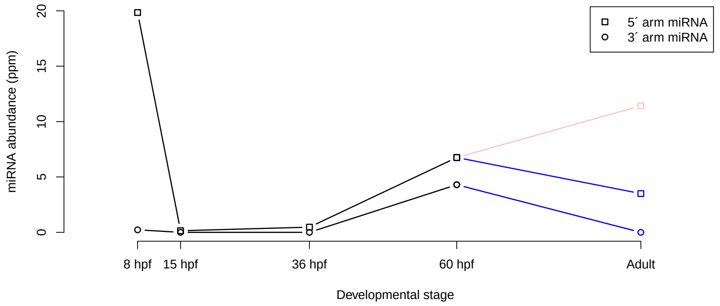 |

(continued on next page)

| Pre-miRNA | miRNA sequences | Abundance profile in development |
|---|---|---|
| bfl-mir-4870 ortholog at Sc0000009 bp 4682265-4682351 (+ strand) | 5´arm: `GAUGUUUGUACUGUCUGUCUGUU`<br>3´arm: `(low abundance)` | 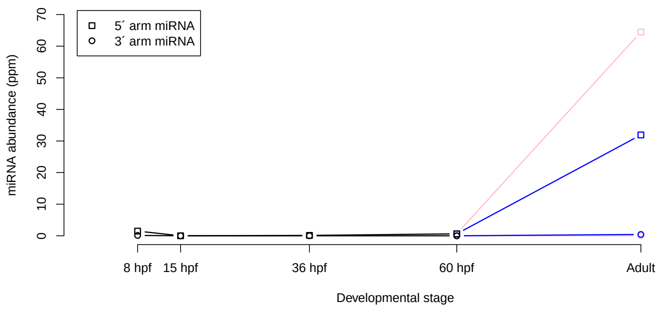 |
| bfl-mir-4866 ortholog at Sc0000043 bp 1850204-1850295 (+ strand) | 5´arm: `UCACACUUGUACUUCUUAGCAU`<br>3´arm: `(low abundance)` | 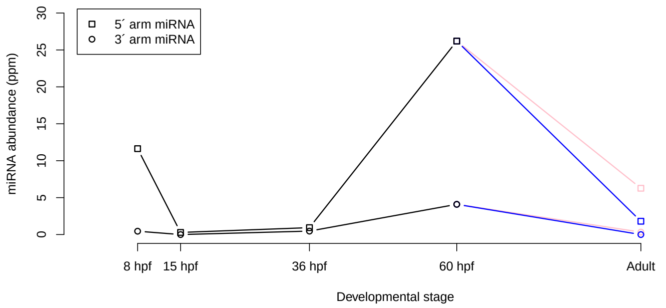 |
| bbe-mir-9 ortholog at Sc0000019 bp 2253983-2254064 (+ strand) | 5´arm: `UCUUUGGUUAUCUAGCUGUAUGA`<br>3´arm: `(low abundance)` | 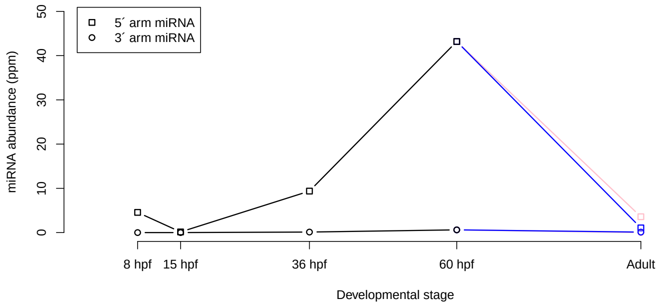 |
| bfl-mir-217 ortholog at Sc0000230 bp 138098-138165 (+ strand) | 5´arm: `UACUGCAUCAGGAACUGAUUGG`<br>3´arm: `AAUCUGUUCCUCAUGCAUGGCU` | 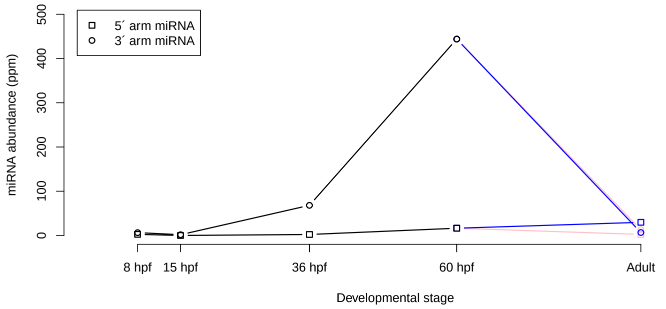 |
